# Structural basis for the midnolin-proteasome pathway and its role in suppressing myeloma

**DOI:** 10.1101/2025.02.22.639686

**Authors:** Christopher Nardone, Jingjing Gao, Hyuk-Soo Seo, Julian Mintseris, Lucy Ort, Matthew C. J. Yip, Milen Negasi, Anna K. Besschetnova, Nolan Kamitaki, Steven P. Gygi, Sirano Dhe-Paganon, Nikhil Munshi, Mariateresa Fulciniti, Michael E. Greenberg, Sichen Shao, Stephen J. Elledge, Xin Gu

## Abstract

The midnolin-proteasome pathway degrades many nuclear proteins without ubiquitination, but how it operates mechanistically remains unclear. Here, we present structures of the midnolin-proteasome complex, revealing how established proteasomal components are repurposed to enable a unique form of proteolysis. While the proteasomal subunit PSMD2/Rpn1 binds to ubiquitinated or ubiquitin-like proteins, we discover that it also interacts with the midnolin nuclear localization sequence, elucidating how midnolin’s activity is confined to the nucleus. Likewise, PSMD14/Rpn11, an enzyme that normally cleaves ubiquitin chains, surprisingly functions non-enzymatically as a receptor for the midnolin ubiquitin-like (Ubl) domain, positioning the substrate-binding Catch domain directly above the proteasomal entry site to guide substrates into the proteasome. Moreover, we demonstrate that midnolin downregulation is critical for the survival of myeloma cells by promoting the expression of its transcription factor substrate IRF4. Our findings uncover the mechanisms underlying the midnolin-proteasome pathway and midnolin downregulation as a driver of multiple myeloma.

## Introduction

Selective protein degradation is essential for maintaining cellular and organismal homeostasis^1^. In eukaryotes, this selectivity is achieved in large part through the action of hundreds of enzymes that covalently attach ubiquitin chains to target proteins^2^, signaling their degradation by the 26S proteasome—a 2.5 MDa protease complex composed of a 19S regulatory particle and a 20S proteolytic core^3^. The 19S regulatory particle canonically recruits ubiquitinated proteins and unfolds them for proteolysis carried out by the 20S core particle. Despite extensive knowledge of this ubiquitin-mediated degradation process, whether and how the 26S proteasome can degrade full-length proteins without ubiquitination has remained a long-standing mystery^4^.

Recently, a parallel pathway to ubiquitin-mediated proteolysis has been discovered in metazoans: the midnolin-proteasome pathway^5^. Midnolin is an inducible protein that associates with the proteasome, either directly or indirectly, to promote the degradation of many nuclear proteins^5,6^. These include transcription factors such as EGR1, Fos, NR4A1, IRF4, NeuroD1, and CBX4, which are all critical for diverse biological processes. For example, EGR1, Fos, and NR4A1 are immediate-early genes that respond to a wide range of cellular signals and regulate cell-type-specific programs involved in growth, differentiation, stress responses, neuronal activity, apoptosis, and inflammation^7,8,9^. CBX4, a component of the polycomb repressive complex (PRC1), acts as a master regulator of transcription during development^10^, while IRF4 is essential for the function of B and T cells^11^, as well as the survival and proliferation of malignant plasma cells in multiple myeloma^12^.

Remarkably, the midnolin-mediated degradation of nuclear substrates by the proteasome occurs independently of ubiquitination and involves three conserved domains of midnolin: an N-terminal ubiquitin-like (Ubl) domain, a “Catch” domain for substrate recognition, and a C-terminal α-helix (αHelix-C) (**Figure S1A**)^5^. However, it remains unknown how midnolin selects nuclear over cytosolic proteins, how midnolin interacts with the proteasome, and how the proteasome, a well-characterized protease for ubiquitinated proteins, enables ubiquitination-independent proteolysis. Furthermore, while midnolin loss is embryonically lethal in mice and flies^6,13^, the significance of the midnolin-proteasome pathway for human health and disease remains to be determined. In this study, we elucidate how midnolin repurposes proteasomal subunits on the 19S regulatory particle to orchestrate ubiquitination-independent proteolysis and uncover a critical role for the pathway in suppressing multiple myeloma.

### Structures of the midnolin-proteasome complex

To determine how midnolin engages the proteasome, we sought to purify the midnolin-proteasome complex for structural analyses. We began by affinity purifying the complex from human embryonic kidney (HEK-293T) cells overexpressing FLAG-tagged midnolin variants, including one N-terminally fused to an EGR1 substrate peptide (**Figures S1B and S1C**). The fused EGR1 peptide may occupy the Catch domain intramolecularly, potentially reducing substrate heterogeneity. Purified samples after size exclusion chromatography (SEC) contained midnolin-proteasome complexes, as indicated by the characteristic Coomassie stain observed after sodium dodecyl sulfate-polyacrylamide gel electrophoresis (SDS-PAGE) (**Figure S1D**) and confirmed by negative stain electron microscopy (**Figure S1E**). These samples were subjected to single-particle cryogenic electron microscopy (cryo-EM) (**Figures S2 and S3**, **Table S1**).

Three-dimensional classification revealed densities associated with the proteasome that could be attributed to midnolin, allowing us to resolve two major conformational states: M1 and M2 (**Figure 1**). In the classical proteasomal mechanism, ubiquitin receptors on the 19S regulatory particle (PSMD2/Rpn1, PSMD4/Rpn10, or ADRM1/Rpn13) recruit ubiquitinated proteins^14-17^, which are then unfolded by translocation through the heterohexameric ATPase ring at the base of the 19S regulatory particle and directly into the 20S core particle for degradation^18^. During the unfolding process, ubiquitin chains on the protein substrate must also be cleaved off by PSMD14/Rpn11, a zinc metalloprotease, to permit successful translocation through the ATPase ring^19-22^. In contrast, midnolin engages the 19S regulatory particle subunits non-canonically, as observed in both the M1 state, characterized by a closed ATPase ring, and the M2 state, in which the ATPase ring is actively translocating a substrate.

**Figure 1.**
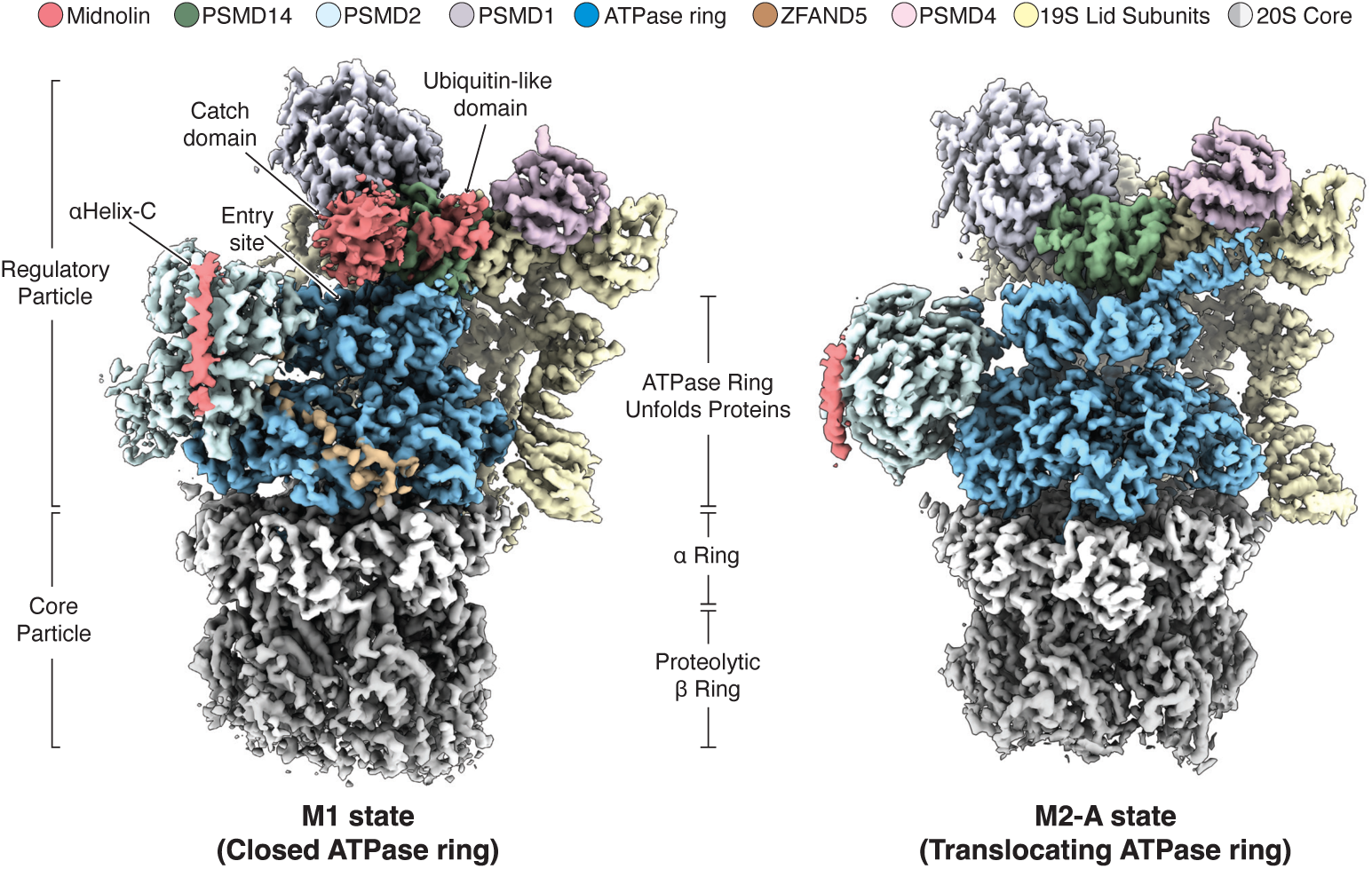
Structures of the midnolin-proteasome complex. Cryo-EM maps of the midnolin-proteasome complex in two distinct conformations. The M1 state displays density for both the midnolin Ubl domain and αHelix-C interacting with the 19S regulatory particle, along with low-resolution density for the Catch domain positioned above the ATPase entry. In contrast, the M2-A state contains density exclusively for αHelix-C.

In the M1 state (3.9 Å), αHelix-C of midnolin binds to PSMD2/Rpn1, and the Ubl domain interacts with PSMD14/Rpn11 (**Figure 1**). We also observed low-resolution density corresponding to the midnolin Catch domain situated directly above the ATPase ring. This configuration would facilitate feeding a Catch-bound substrate into the ATPase for unfolding and subsequent degradation. The overall architecture of the M1 proteasome closely resembles the previously reported Z+D state, which features ZFAND5 bound to the ATPase, PSMD2 positioned near the substrate entry site, and PSMD14 offset from the closed central pore^23^. Thus, the M1 state, showcasing the three functional domains of midnolin, represents a midnolin-proteasome intermediate poised to deliver a substrate for unfolding.

In the EGR1-midnolin fusion dataset, we observed an M1-like state that lacked density for the Ubl and Catch domains. This conformation was abundant in the EGR1-midnolin dataset but was not found in the other datasets, possibly due to steric hindrance from the N-terminal EGR1 peptide fusion (**Figure S2**).

Two M2 states (A: 3.1 Å and B: 3.4 Å) were identified across all datasets. Both states exhibited only the midnolin αHelix-C bound to the regulatory particle but displayed ATPase configurations at different stages of active translocation, with distinguishable density corresponding to a substrate polypeptide (**Figure S4**). The overall architecture of the more abundant M2-A state resembled the previously reported human ED2 or yeast 4D proteasomal states^24,25^. These results suggest that, in the M1 state, the midnolin Ubl domain positions the Catch domain above the ATPase ring for substrate delivery, while in the M2 state, the Ubl domain disengages from PSMD14/Rpn11 during protein degradation.

### Mechanism for assembling the midnolin-proteasome complex in the nucleus

The M1 and M2 states showed a binding interface between midnolin αHelix-C and a region of PSMD2 that we refer to as the ‘M site’ (**Figure 2A)**. The M site is distinct from the ubiquitin and Rad23 binding surfaces (T1 and NT) but partially overlaps with the USP14 binding site (T2)^17,26-28^. Surprisingly, the arginine-rich region of αHelix-C (R405-R413), which serves as a nuclear localization sequence (NLS)^5^, formed electrostatic interactions with acidic residues of PSMD2 (E315, D316, and E319) (**Figure 2A**). The use of the midnolin NLS to bind the proteasome is likely conserved throughout evolution as predicted by AlphaFold-multimer^29^ (**Figure S5**). We validated the structural models by mutagenizing the M site of PSMD2 and found that E319, L426, Y434, L465, and Y469 of PSMD2 were required to bind midnolin (**Figure S6**). E319 engages the midnolin NLS electrostatically, while L426, Y434, L465, and Y469 engage the N-terminal half of αHelix-C primarily through hydrophobic interactions. Thus, midnolin associates stably with the proteasome in part using an NLS.

**Figure 2.**
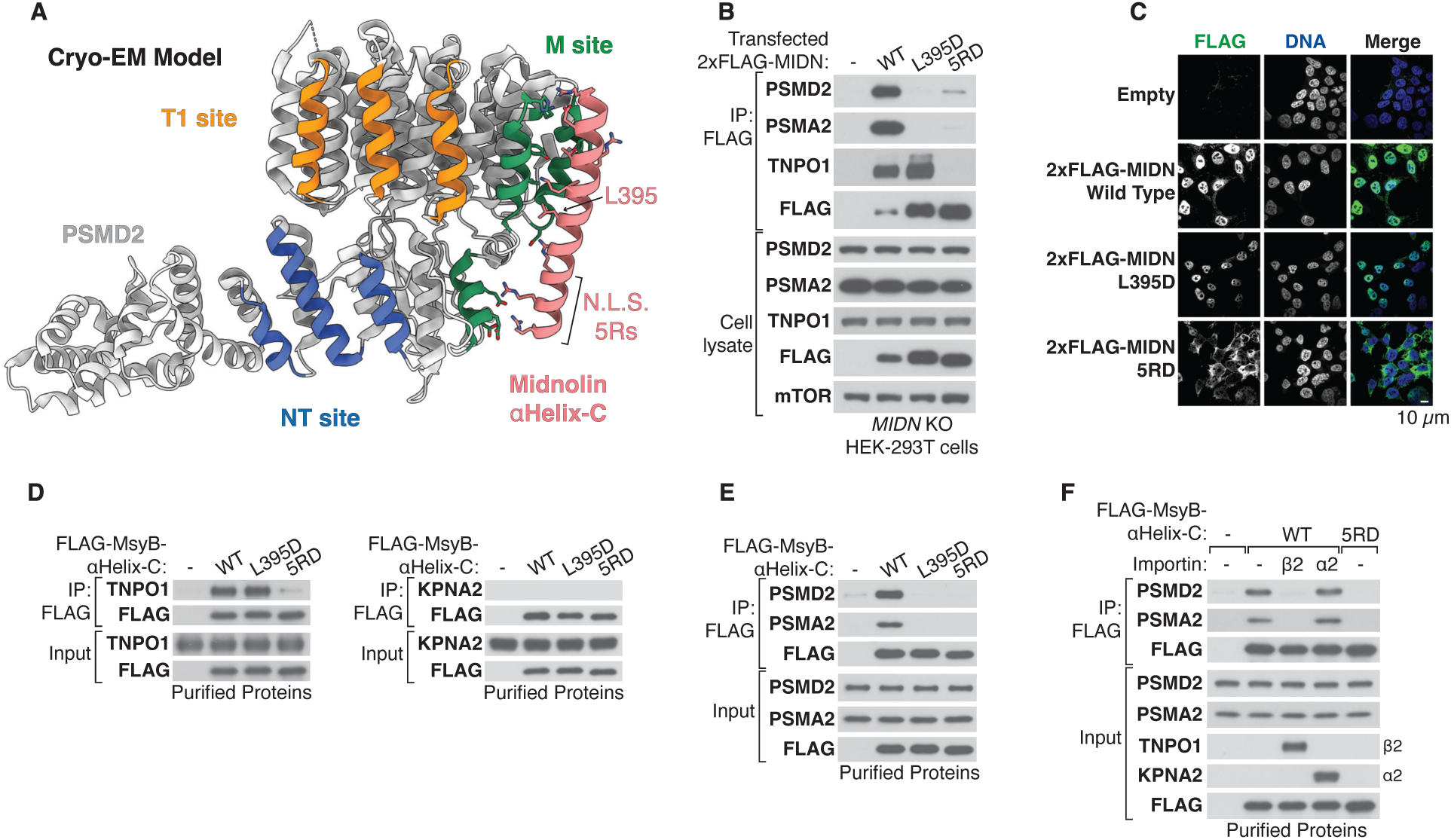
Mechanism for assembling the midnolin-proteasome complex in the nucleus. (A) Atomic model of midnolin αHelix-C in complex with PSMD2/Rpn1, highlighting the role of the midnolin nuclear localization sequence (NLS) in mediating the interaction. (B) Immunoblotting of anti-FLAG immunoprecipitants from *MIDN* knockout HEK-293T cells transiently overexpressing 2xFLAG-midnolin using a CMV promoter. (C) Anti-FLAG immunofluorescence of *MIDN* knockout HEK-293T cells stably expressing 2xFLAG-midnolin. Cells were treated with 10 µM MG132 for 4 hours. (D) *In vitro* co-immunoprecipitation followed by immunoblotting. Purified FLAG-MsyB-αHelix-C were immobilized onto anti-FLAG beads and incubated with pure TNPO1 or KPNA2, (E) human proteasomes, or (F) human proteasomes with either TNPO1 or KPNA2.

The proteasome is localized in both the cytosol and nucleus^30^. We hypothesized that if the midnolin-proteasome complex assembled in the cytosol, it could lead to unproductive degradation of nuclear proteins before they can enter the nucleus to perform their functions, as well as the degradation of cytosolic proteins that are not meant to be targeted by the midnolin-proteasome pathway. The observation that the NLS is critical for proteasome binding suggested a mechanism for assembling the midnolin-proteasome complex primarily in the nucleus. Nuclear import receptors may prevent midnolin from associating with the cytosolic proteasome by sterically occluding αHelix-C, facilitating the assembly of the midnolin-proteasome complex primarily in the nucleus.

To investigate this mechanism, we generated mutants in αHelix-C and tested their impact on proteasome binding and nuclear localization. Mutation of leucine 395 (L395D) or five arginines comprising the NLS (5RD) abolished the midnolin-proteasome interaction (**Figure 2B**). Although wild-type and L395D midnolin were localized primarily within the nucleus, 5RD midnolin was cytosolic (**Figure 2C**). Thus, midnolin does not require proteasome binding to enter the nucleus.

When fused to maltose-binding protein (MBP), wild-type αHelix-C of midnolin, but not the L395D or 5RD variants, bound to the proteasome (**Figure S7A**). Both wild-type and L395D αHelix-C promoted the nuclear localization of MBP, while 5RD αHelix-C remained cytosolic (**Figure S7B**). To identify potential mechanisms regulating the nuclear transport of midnolin, we immunoprecipitated MBP-αHelix-C variants and conducted mass spectrometry. Among the most enriched proteins were members of the importin β family (TNPO1, TNPO2, KPNB1, and IPO5) (**Figure S7C, Table S2**). The importin β family recognizes a positively charged NLS and transports cargo directly into the nucleus through the nuclear pore complex^31-34^. AlphaFold3^35^ predicted a direct interaction between the midnolin NLS and TNPO1, which we validated with single amino acid substitutions in αHelix-C (**Figure S7D-S7F**). However, midnolin mutants that fail to bind TNPO1 still localize to the nucleus (**Figure S7G**), likely because these variants still bind to other importins (**Figure S7F**). Therefore, midnolin αHelix-C is necessary and sufficient for both nuclear transport, which occurs via redundant importin β family members, and for binding to the proteasome through PSMD2.

To investigate whether αHelix-C binds directly to TNPO1 and the proteasome in a mutually exclusive manner, we reconstituted the interactions using purified proteins (**Figure S8**). Both wild-type and L395D αHelix-C bound to TNPO1 but not to KPNA2, an unrelated nuclear import receptor of the importin α family (**Figure 2D**). Only wild-type αHelix-C associated with the proteasome (**Figure 2E**). We then immunoprecipitated αHelix-C in the presence of the proteasome and either TNPO1 or KPNA2. The interaction between the proteasome and αHelix-C was disrupted in the presence of TNPO1 (β2), but not with KPNA2 (α2) (**Figure 2F**). Furthermore, αHelix-C bound to TNPO1 with approximately 50 times greater affinity than to PSMD2, as measured by biolayer interferometry (**Figure S8E**). Thus, TNPO1 and the proteasome compete to bind the NLS-containing αHelix-C. These findings suggest that this competition between nuclear import receptors and the proteasome restricts the activity of midnolin to the nucleus.

### PSMD14/Rpn11 acts as a Ubl receptor to position the Catch domain above the proteasomal entry site

Although the Ubl domain is required for midnolin’s function, the precise mechanism by which it promotes ubiquitination-independent degradation remains unclear. The M1 state revealed that the Ubl domain of midnolin binds to the crucial deubiquitinase PSMD14/Rpn11 (**Figure 1**), an interaction likely conserved across evolution (**Figure S9**). Notably, the Ubl domain is followed by a short linker positioned near the active site of PSMD14, resembling the structure of a ubiquitin chain poised for cleavage from ubiquitinated proteins during translocation^24^ (**Figure 3A**).

**Figure 3.**
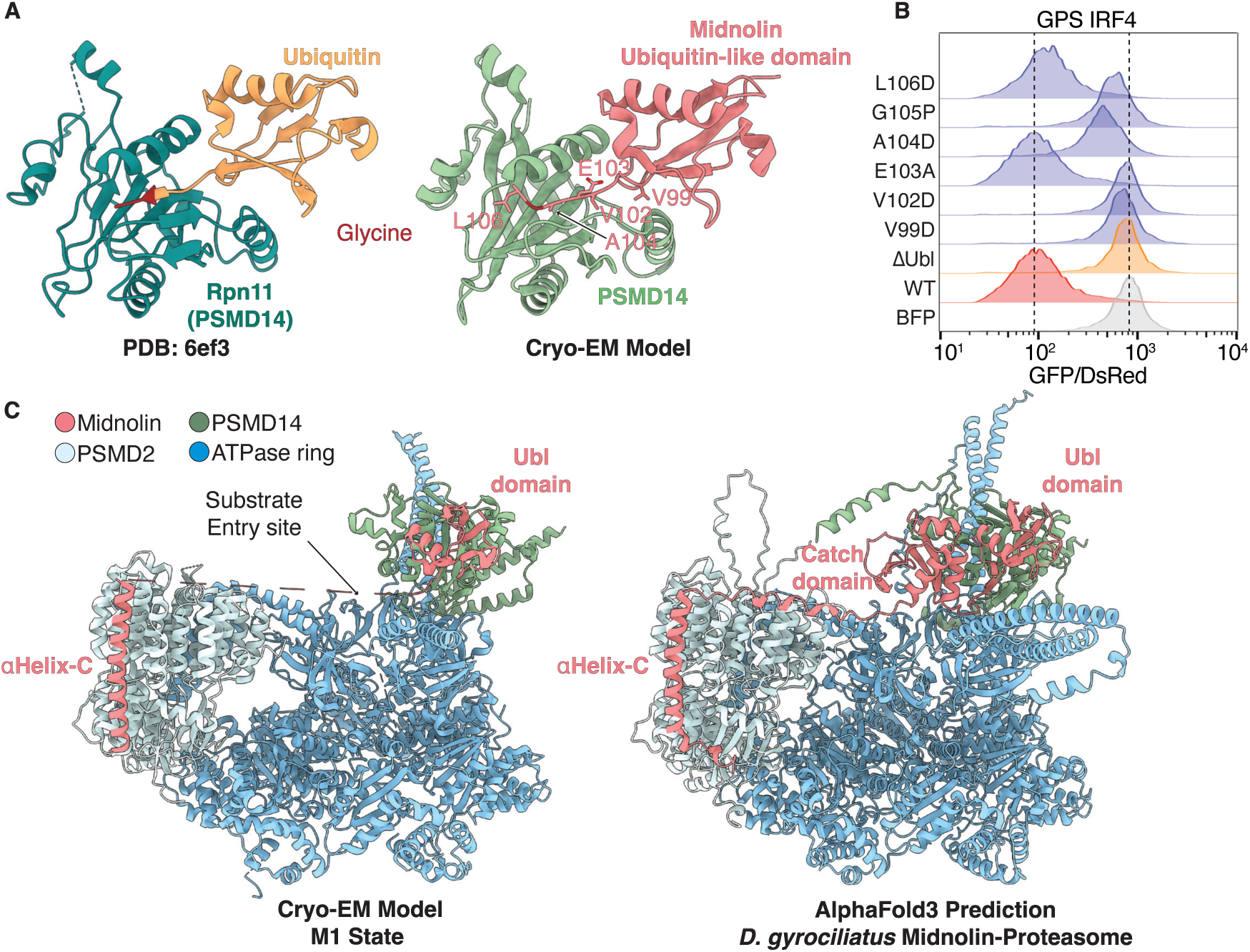
PSMD14/Rpn11 acts as a Ubl receptor to position the Catch domain above the proteasomal entry site. (A) Atomic model of the midnolin ubiquitin-like domain bound to the deubiquitinase PSMD14/Rpn11. For comparison, the structure of ubiquitin bound to Rpn11 is shown (PDB: 6EF3)^24^. (B) *MIDN* knockout HEK-293T cells stably expressing a dual-fluorescence IRF4 stability reporter were transfected with either control BFP or wild-type and Ubl-mutant midnolin co-expressing BFP using an EF-1α promoter. The GFP/DsRed ratio of BFP+ cells (∼10,000 cells) was analyzed by flow cytometry two days post-transfection. (C) AlphaFold3 prediction of *D. gyrociliatus* midnolin-PSMD2-PSMD14-ATPase (right) closely resembles the cryo-EM model of human M1 midnolin-proteasome complex (left).

To test the requirement of the Ubl-PSMD14 interaction for substrate degradation, we employed a Global Protein Stability (GPS) reporter^36,37^ of the midnolin substrate IRF4. This dual fluorescent reporter expresses DsRed as an internal control and green fluorescent protein (GFP) fused to IRF4 from the same bicistronic mRNA. The GFP/DsRed ratio, measured by flow cytometry, provides the relative stability of the fusion protein. We overexpressed wild-type and mutant versions of midnolin in *MIDN* knockout HEK-293T cells stably expressing the IRF4 reporter. Mutagenesis of the linker residues buried within PSMD14 (V99, V102, A104) abolished midnolin function, but disrupting the solvent-exposed linker residues (E103 and L106) had little to no effect (**Figure 3B**). To validate the interaction between midnolin and PSMD14 further, we performed crosslinking mass spectrometry using purified midnolin-proteasome complexes. Analysis revealed a unique crosslink between PSMD14 and the midnolin Ubl domain, consistent with the cryo-EM model (**Figure S10A, Table S3**). Thus, PSMD14 acts as a receptor for the midnolin Ubl domain.

The Catch domain is separated from the Ubl domain by a short linker sequence (12 amino acids) and is clearly visible in our density map of the M1 proteasomal state. This Catch density is positioned between the Ubl domain and αHelix-C, directly above the substrate entry site of the ATPase ring (**Figure 1**). However, the Catch domain appears at low resolution, likely due to its flexibility. AlphaFold3 predictions using the ATPase subunits, PSMD2, PSMD14, and midnolin confirm the position of the Catch domain above the substrate entry site (**Figure S10B**).

Notably, we identified a minimal midnolin from the segmented worm *Dimorphilus gyrociliatus*, which contains only the Ubl domain, Catch domain, and αHelix-C, with minimal unstructured regions in between (**Figure S11A**). This worm species has a highly compacted genome^38^. When expressed in human cells, *D. gyrociliatus* midnolin localized to the nucleus, bound the proteasome, and promoted the degradation of the EGR1 substrate (**Figure S11B-S11D**). AlphaFold3 also predicted that *D. gyrociliatus* midnolin interacts with the proteasome in a manner identical to human midnolin (**Figure 3C**).

Importantly, the Ubl and αHelix-C domains of this minimal midnolin can only simultaneously occupy the M1 proteasomal state. In other reported proteasomal states, the distance between PSMD14 (Ubl) and PSMD2 (αHelix-C) is too large. These findings suggest that the three functional domains of midnolin—Ubl, Catch, and αHelix-C—are both necessary and sufficient for its function. Furthermore, the M1 proteasomal state we identified is essential for enabling the coordinated action of these domains in mediating substrate degradation.

### The Catch domain can enhance the substrate interaction by forming an FG zipper

The midnolin Catch domain selects substrates^5^. AlphaFold suggests that the Catch domain consists of two discontinuous but semi-symmetrical regions, each comprising two β strands and two or three α helices^39^. Using AlphaFold-multimer, we previously found that an unstructured region within a midnolin substrate may adopt a β strand upon binding to the Catch domain, completing an anti-parallel β sheet tertiary structure^5^. However, by cryo-EM we could not observe the Catch domain with sufficient resolution to confirm the AlphaFold prediction. Therefore, we solved a crystal structure of the Catch domain bound to the EGR1 substrate peptide at a resolution of 2.5 Å (**Figure 4A**).

**Figure 4.**
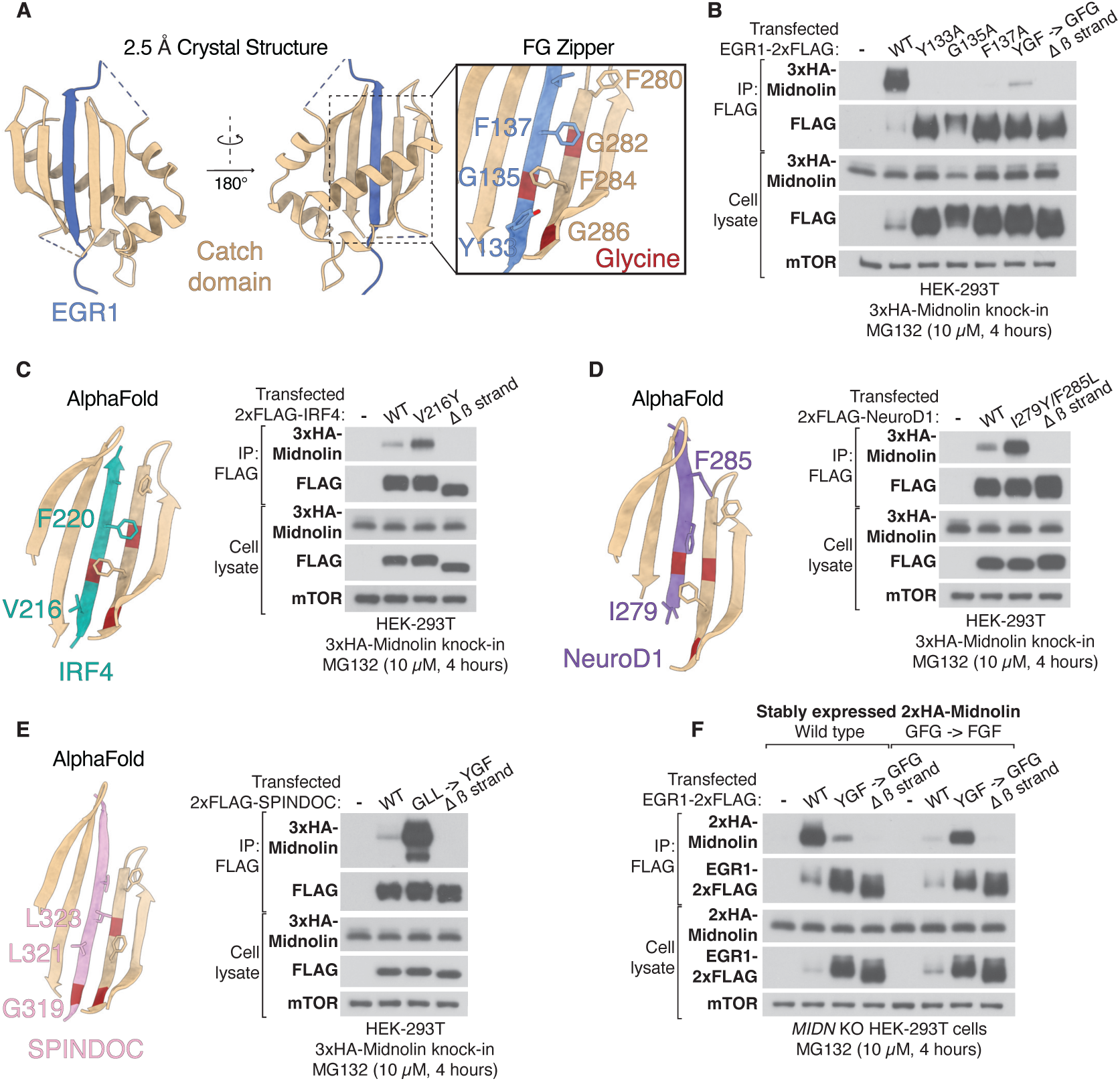
The Catch domain can enhance the substrate interaction by forming an FG zipper. (A) Crystal structure of the EGR1-Catch fusion protein at 2.5 Å resolution. The interaction between EGR1 and the Catch domain is facilitated by alternating phenylalanine-glycine residues, forming an FG zipper. (B) Immunoblotting of anti-FLAG immunoprecipitates from HEK-293T cells expressing endogenous 3xHA-midnolin and transiently overexpressing EGR1-2xFLAG via a CMV promoter. Cells were treated with 10 µM MG132 for 4 hours. (C) AlphaFold-multimer prediction of the IRF4-midnolin complex shows an incomplete FG zipper, with valine 216 replacing tyrosine. The same immunoblot assay as in (b) was performed using cells transfected with 2xFLAG-IRF4. (D) AlphaFold-multimer prediction of the NeuroD1-midnolin interaction reveals an incomplete FG zipper, with isoleucine 279 replacing tyrosine and phenylalanine 285 sterically clashing with phenylalanine 280 of midnolin. The same assay as in (b) was conducted with cells transfected with 2xFLAG-NeuroD1. (E) AlphaFold-multimer prediction of SPINDOC-midnolin shows a missing FG zipper, with glycine 319, leucine 321, and leucine 323 substituting for tyrosine, glycine, and phenylalanine, respectively. The same assay as in (b) with cells transfected with 2xFLAG-SPINDOC. (F) The same assay as in (b) was performed using *MIDN* knockout HEK-293T cells reconstituted with either wild-type or zipper-swapped 2xHA-midnolin from a CMV promoter using lentivirus.

The substrate-free Catch domain, like full-length midnolin, proved very challenging to express and purify from bacteria. However, fusing the EGR1 peptide to the Catch domain significantly enhanced bacterial expression (**Figure S12A)**. This enabled us to purify proteins that formed well-defined crystals (**Figure S12B and S12C**). The crystal structure confirmed the β strand complementation mechanism. Among all proteins identified as midnolin substrates, EGR1 bound most effectively to the Catch domain, as demonstrated using co-immunoprecipitation experiments^5^. To understand the biochemical basis for this observation, we examined the residues of EGR1 and midnolin buried within the Catch domain core. We identified alternating phenylalanine and glycine residues from both midnolin and EGR1, which were well-supported by the density (**Figure S12D)** and highly conserved throughout evolution (**Figure S13**). This phenylalanine-glycine (FG) zipper could explain how EGR1 binds to midnolin so strongly.

To test the necessity of the FG zipper, we introduced mutations within the EGR1 β strand and performed a co-immunoprecipitation experiment from cells endogenously expressing epitope-tagged midnolin. The midnolin-EGR1 interaction was disrupted by substituting the FG zipper residues with alanine or changing the sequence from YxGxF to GxFxG (**Figure 4B**).

We revisited our previous AlphaFold-multimer predictions for midnolin-substrate interactions^5^. IRF4 is missing one aromatic residue to complete the FG zipper. Substituting valine 216 with tyrosine enhanced the midnolin-IRF4 binding (**Figure 4C**). NeuroD1 not only lacks one aromatic residue to complete the FG zipper but also contains phenylalanine 285, which may sterically clash with phenylalanine 280 of midnolin. Replacing isoleucine 279 with tyrosine and phenylalanine 285 with leucine significantly improved the midnolin-NeuroD1 binding (**Figure 4D**). Finally, for SPINDOC, which lacks an FG zipper, substituting glycine 319, leucine 321, and leucine 323 to tyrosine, glycine, and phenylalanine, respectively, dramatically enhanced the midnolin-SPINDOC binding (**Figure 4E**). Thus, the interaction between midnolin and its high-affinity substrates is driven not only by β strand complementation but also by specific residues that can form a partial or complete FG zipper with the Catch domain.

To reinforce the importance of the FG zipper, we swapped the zipper residues between midnolin and EGR1. Specifically, EGR1 was modified from YxGxF to GxFxG, while the Catch domain was altered from GxFxG to FxGxF. The wild-type EGR1 predominantly bound to the wild-type midnolin, while the swapped EGR1 primarily interacted with the swapped midnolin (**Figure 4F**). This result suggests that the midnolin Catch domain can be engineered to bind other proteins of interest that do not interact with the wild-type Catch domain.

### Midnolin is downregulated in multiple myeloma and suppresses cancer cell growth via IRF4 degradation

Among various transcription factors, IRF4 stands out as one of the most efficiently degraded substrates of the midnolin-proteasome pathway^5^. IRF4 is critical for establishing plasma cell identity and plays a pivotal role in the pathogenesis of multiple myeloma (MM)^12^. MM is an aggressive hematologic cancer characterized by the clonal expansion of malignant plasma cells in the bone marrow. This uncontrolled proliferation leads to complications such as bone damage, anemia, abnormal blood clotting due to thrombocytopenia, and immune suppression caused by a reduction in white blood cells^40^. Additionally, the overproduction of antibodies in MM can result in kidney and organ damage. Reducing IRF4 expression by as little as 50% significantly impairs myeloma cell growth^12^, and genetic dependency analyses have identified IRF4 as a critical vulnerability across all MM cell lines^41^. Given that IRF4 is one of the most efficiently degraded proteins via the midnolin-proteasome pathway, we hypothesize that midnolin may play a significant role in MM progression.

To test this hypothesis, we first examined midnolin transcript levels across various cancer cell lines^42^. Midnolin mRNA was selectively downregulated approximately 4- to 5-fold in multiple myeloma (MM) cell lines compared to those derived from solid tumors, lymphomas, or leukemias (**Figure 5A**). This downregulation was further confirmed via quantitative polymerase chain reaction (qPCR) in different lymphoma and myeloma cell lines (**Figure S14A**). Additionally, midnolin mRNA levels were reduced in an RNA sequencing dataset^43^ of CD138+ primary plasma cells from newly diagnosed MM patients compared to healthy controls (**Figure 5B**). We also observed midnolin downregulation in premalignant conditions, such as monoclonal gammopathy of undetermined significance (MGUS) and smoldering multiple myeloma (SMM) (**Figure 5C**). These findings suggest that midnolin downregulation is an early and recurring event in MM progression.

**Figure 5.**
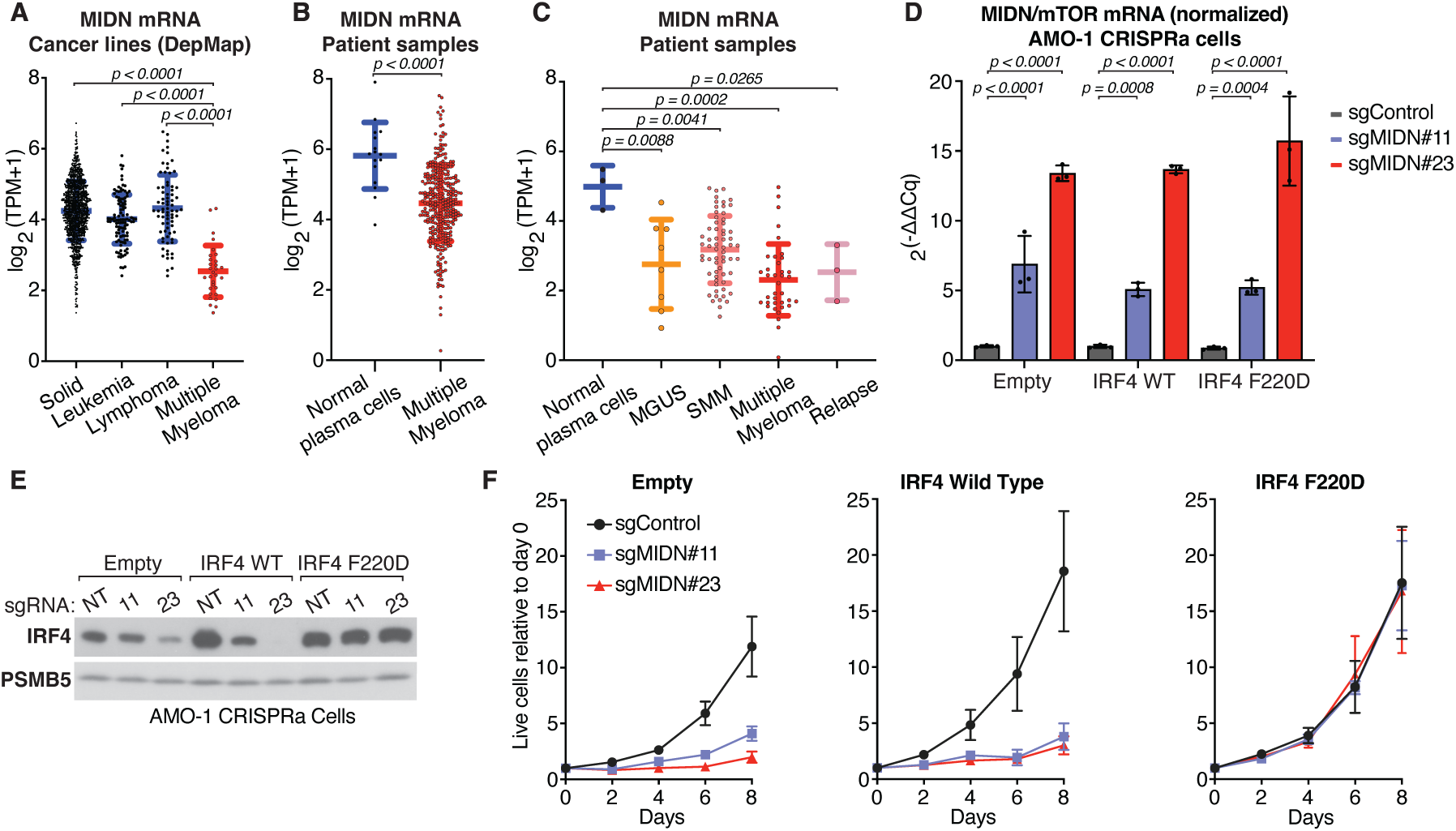
Midnolin is downregulated in multiple myeloma and suppresses cancer cell growth via IRF4 degradation. (A) RNA sequencing data from various cancer cell lines, including solid tumors (n=1257) and three hematological malignancies (leukemia n=105; lymphoma n=74; multiple myeloma n=30)^42^. (B) RNA sequencing of primary plasma cells isolated from healthy donors (n=23) and newly diagnosed multiple myeloma patients (n=507)^43^. (C) RNA sequencing of primary plasma cells from healthy donors (n=3), patients with pre-malignant conditions, including monoclonal gammopathy of undetermined significance (MGUS, n=8) and smoldering multiple myeloma (SMM, n=65), as well as patients with newly diagnosed multiple myeloma (n=37) and relapsed multiple myeloma (n=3). (D) AMO-1 cells were engineered to express either an empty vector, wild-type IRF4, or the IRF4 (F220D) mutant, which cannot bind to midnolin. These AMO-1 cells also express nuclease-deactivated Cas9-VPR and either control or MIDN-targeting sgRNAs to induce midnolin expression. The qPCR data (n=3) illustrate the relative expression levels of MIDN compared to mTOR transcripts in these cell lines. (E) Immunoblotting of the same AMO-1 cell lines as in (d). (F) Proliferation of the cell lines described in (d) was assessed using the Cell Titer Glow (CTG) assay to measure live cells. In (a-d) and (f), the values are shown as the mean ± the standard deviation. In (d) and (f), the data is of biological replicates. Statistical analysis was performed using Kruskal-Wallis test (a), two-tailed unpaired *t*-test (b), two-way ANOVA followed by Tukey’s multiple comparisons test (c, d, and f). In (f), the p values shown represent comparisons at day 8.

To assess the functional consequences of midnolin downregulation, we restored endogenous midnolin expression in MM cells using CRISPR-activation. AMO-1 cells were transduced with lentivirus to stably overexpress nuclease-deactivated Cas9 fused to three transcriptional activators (VP64, p65, and Rta), along with either a non-targeting single-guide RNA control or sgRNAs targeting the *MIDN* locus. Introducing the midnolin-targeting sgRNAs resulted in increased midnolin transcripts (**Figure 5D**) and decreased IRF4 protein levels (**Figure 5E and S14B**). Restoring midnolin expression to levels comparable to healthy plasma cells (sgRNA #11) inhibited cell growth over time compared to the non-targeting control (**Figure 5F**). Remarkably, the proliferation defect was rescued entirely by expressing an FG-zipper mutant IRF4 (F220D) that cannot bind to the Catch domain (**Figure 5F**). Together, these results indicate that midnolin suppresses multiple myeloma by promoting IRF4 degradation.

## Discussion

Midnolin regulates the proteasomal degradation of many key transcription factors, yet the molecular events that mediate this ubiquitination-independent process were unclear. In this study, we elucidate the underlying mechanisms of the midnolin-proteasome pathway and propose the following model (**Fig. 6**): after midnolin is synthesized, its NLS-containing αHelix-C is bound by the importin β family of nuclear import receptors, which act as gatekeepers to prevent the midnolin-proteasome interaction in the cytosol. Upon nuclear translocation, midnolin is released from the import receptor and assembles with the proteasome through PSMD2/Rpn1. Unstructured regions of substrates form a β strand to bind the Catch domain. This interaction can be enhanced by an FG zipper, in which alternating phenylalanine-glycine residues in both binding partners interact. The midnolin Ubl domain then binds PSMD14/Rpn11, which is typically required to cleave the ubiquitin chain. Here, however, PSMD14/Rpn11 acts as a Ubl receptor to position the Catch domain directly above the ATPase ring, directing a bound substrate into the proteasome for degradation. The similarity in how the midnolin Ubl domain and ubiquitin engage PSMD14/Rpn11 clarifies how midnolin initiates proteasomal degradation of bound substrates independently of ubiquitination.

**Figure 6.**
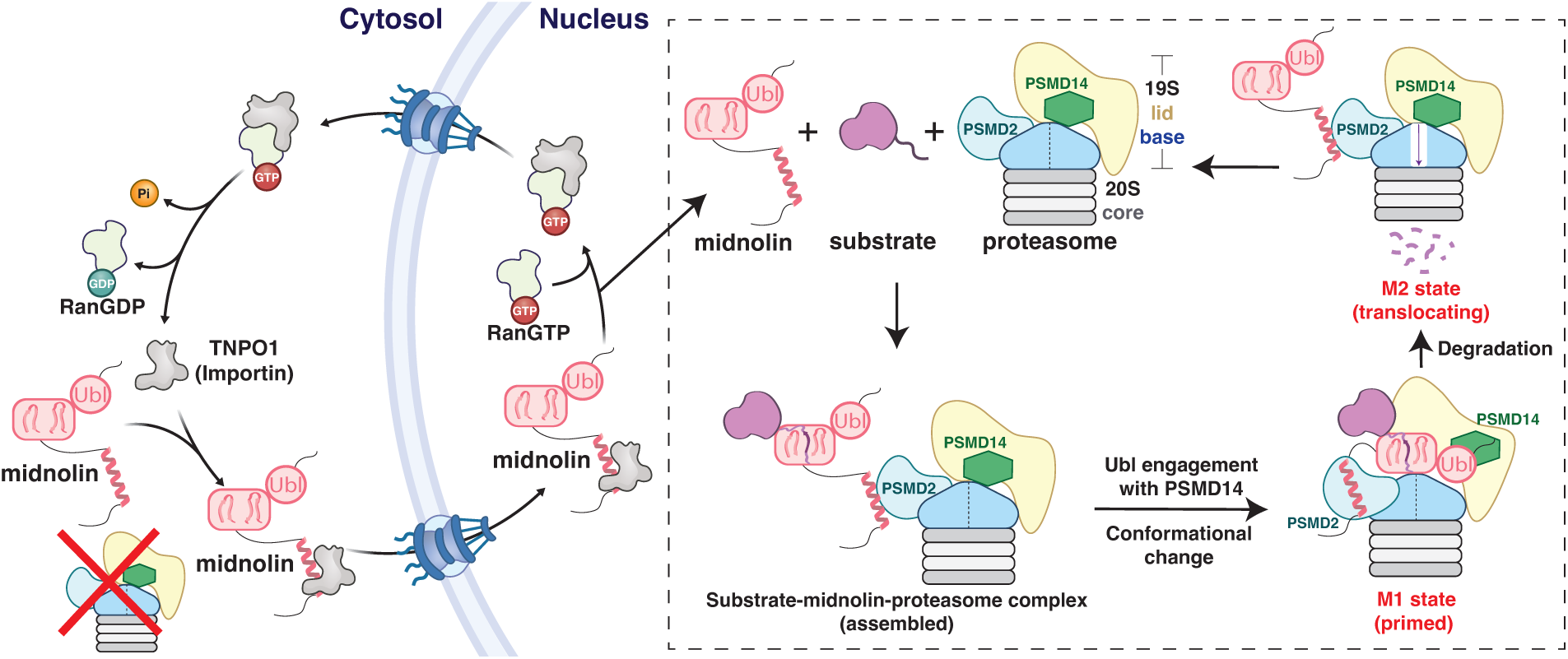
Model for how the midnolin-proteasome pathway degrades nuclear proteins independently of ubiquitination. The importin β family of nuclear import receptors bind to midnolin upon its synthesis, preventing proteasomal association in the cytosol and facilitating nuclear transport. The high concentration of Ran-GTP in the nucleus releases midnolin, enabling its interaction with the 19S regulatory particle via PSMD2/Rpn1. The midnolin ubiquitin-like domain (Ubl) binds to the proteasomal component PSMD14/Rpn11, which positions the Catch domain above the ATPase ring to initiate substrate degradation (M1 state). During translocation (M2 state), the midnolin Ubl domain may disengage from PSMD14/Rpn11. After substrate translocation, midnolin may be reused to deliver other substrates or degraded due to proximity to the ATPase ring. Created with BioRender.com.

Our dissection of how midnolin engages the proteasome to enable ubiquitination-independent proteolysis reveals new functions for proteasomal components and introduces new concepts in protein regulation with broader implications. For instance, the dual role of midnolin’s nuclear localization sequence in regulating both its localization and interaction with the proteasome represents an ingenious way to restrict the activity of a protein to the nucleus. Other proteins with binding partners in both the cytosol and nucleus might employ a similar strategy to achieve location-specific functions.

Moreover, whether midnolin is unique or other proteins with similar characteristics could promote ubiquitination-independent proteolysis remains unknown. Notably, hundreds of proteins across evolutionary kingdoms have been identified to contain at least one ubiquitin-like (Ubl) domain^44^. While the mechanism of action of the Ubl domain for some Ubl-containing proteins is known, most remain unclear and could employ at least some of the mechanisms used by midnolin that we described here. For example, positioned strategically above the ATPase entry, PSMD14/Rpn11 is ideally situated to recruit proteins containing a Ubl domain, which could either be directly degraded by the proteasome or, like midnolin, act to deliver bound cargo for degradation. While PSMD14/Rpn11 has traditionally been known for its role in processing ubiquitin chains, our findings suggest it may also serve as a general receptor for ubiquitination-independent proteolysis.

Although midnolin is essential for embryonic development in mice and flies, its physiological and pathological roles remain unclear. Midnolin has been suggested to regulate development, sleep, metabolism, and immune cell function^6,13,45,46^. Given its broad range of substrates, midnolin is likely involved in additional, yet-undiscovered cellular processes. In this study, we found that midnolin is selectively downregulated in primary malignant plasma cells from patients with multiple myeloma compared to healthy donors. We also discovered that restoring midnolin expression in myeloma cancer cells is sufficient to suppress their growth by degrading a single transcription factor, IRF4. Remarkably, a point mutation in IRF4 that blocks its interaction with midnolin completely rescues the growth suppression of myeloma cells caused by midnolin upregulation. These findings unveil a previously unrecognized regulatory pathway in multiple myeloma and raise important questions for future study.

One such question concerns the specific mechanisms responsible for the downregulation of midnolin in MM. The downregulation could potentially be driven by DNA hypermethylation of the *MIDN* locus, a misregulation of transcription factors controlling *MIDN* expression, or a reduction in the stability of *MIDN* transcripts. Additionally, we observed midnolin downregulation in pre-malignant plasma cells, indicating that midnolin loss occurs early during disease progression. Thus, another crucial question is whether the downregulation of midnolin in healthy plasma cells is necessary and/or sufficient for the progression of MM in animal models. Despite recent advances in therapy, MM remains largely incurable, with nearly all patients eventually relapsing due to treatment resistance^47^. A deeper understanding of the role of midnolin in MM may open new avenues for therapy, particularly if methods can be developed to enhance midnolin expression or activity.

## STAR METHODS

### KEY RESOURCES TABLE

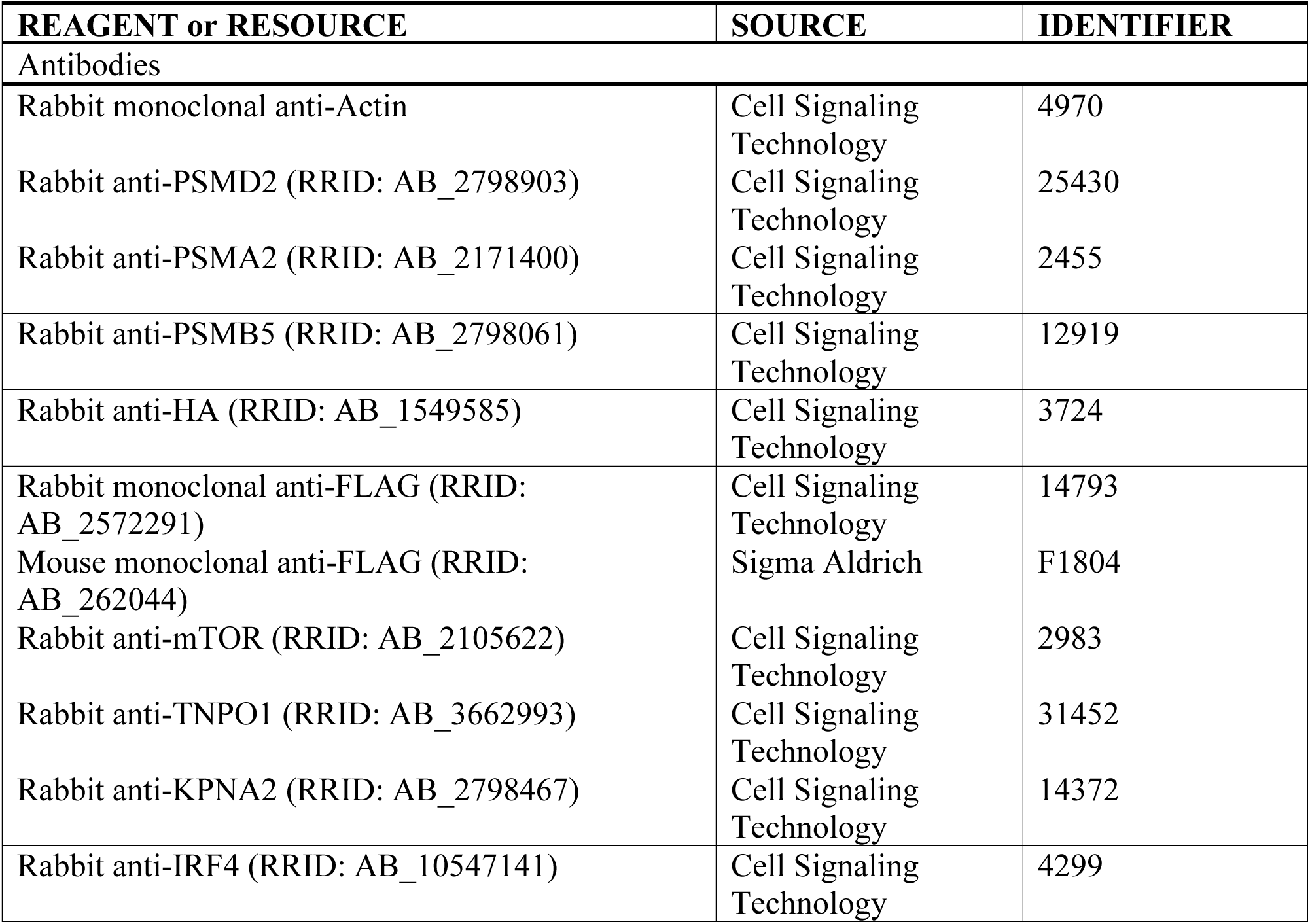
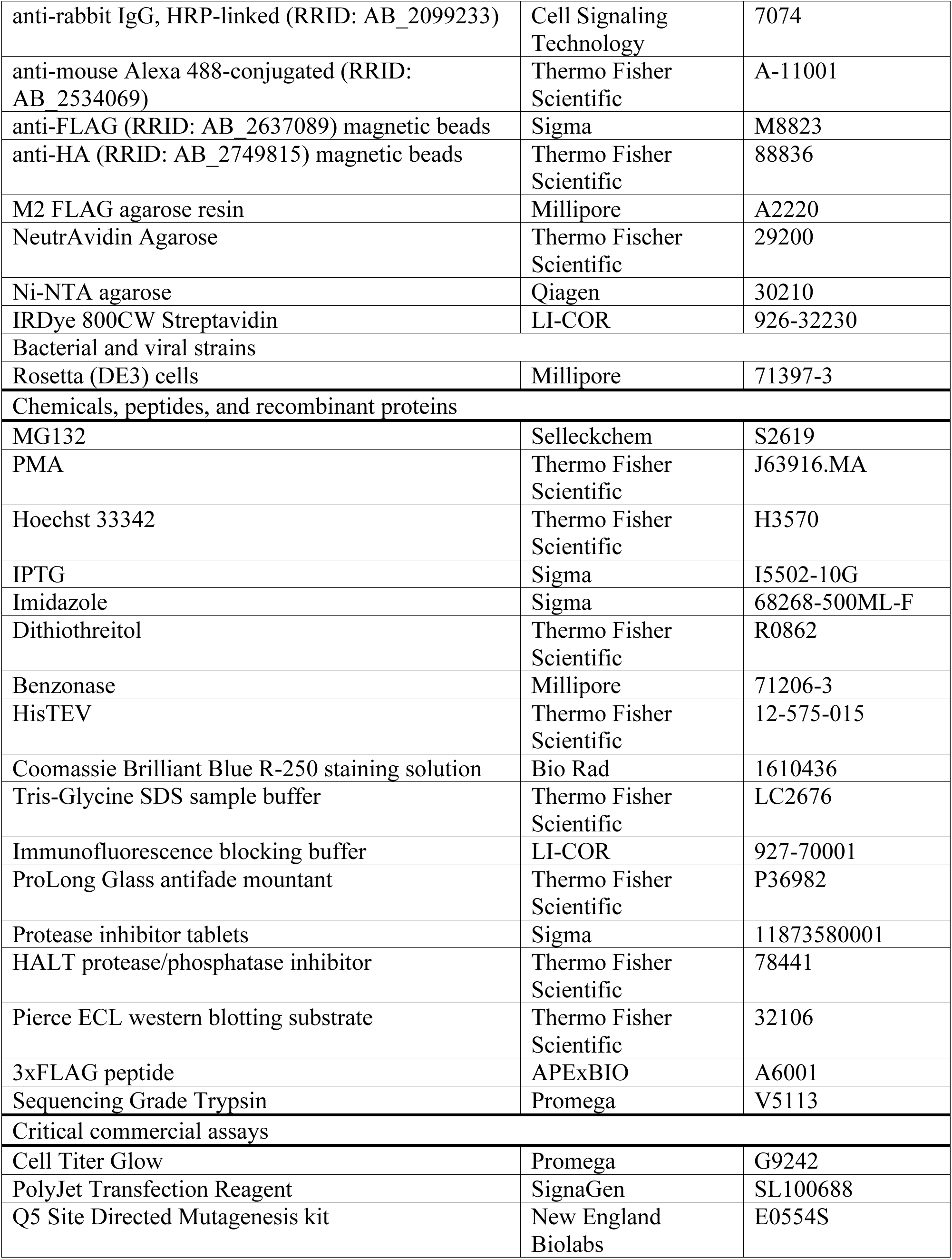
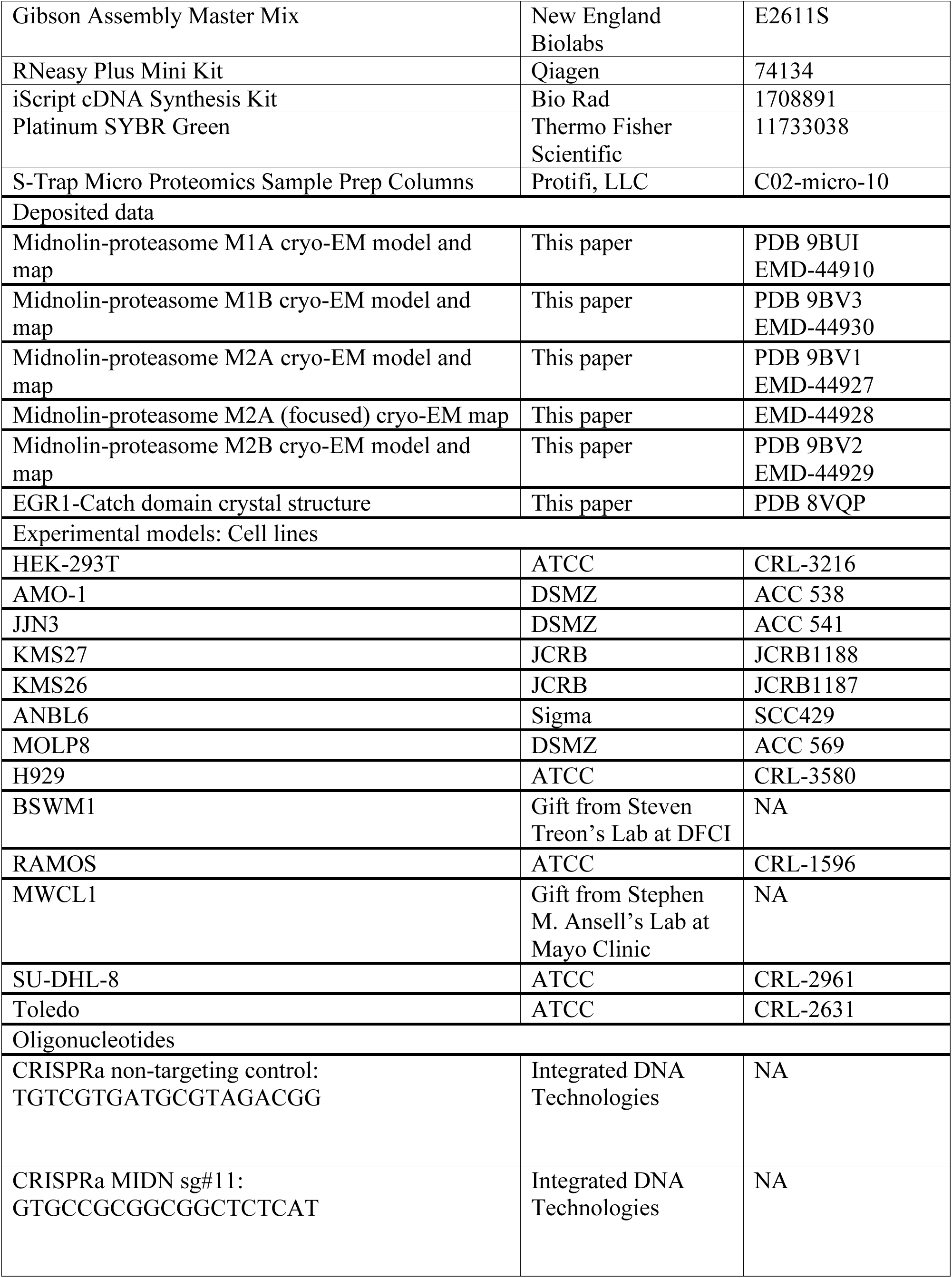
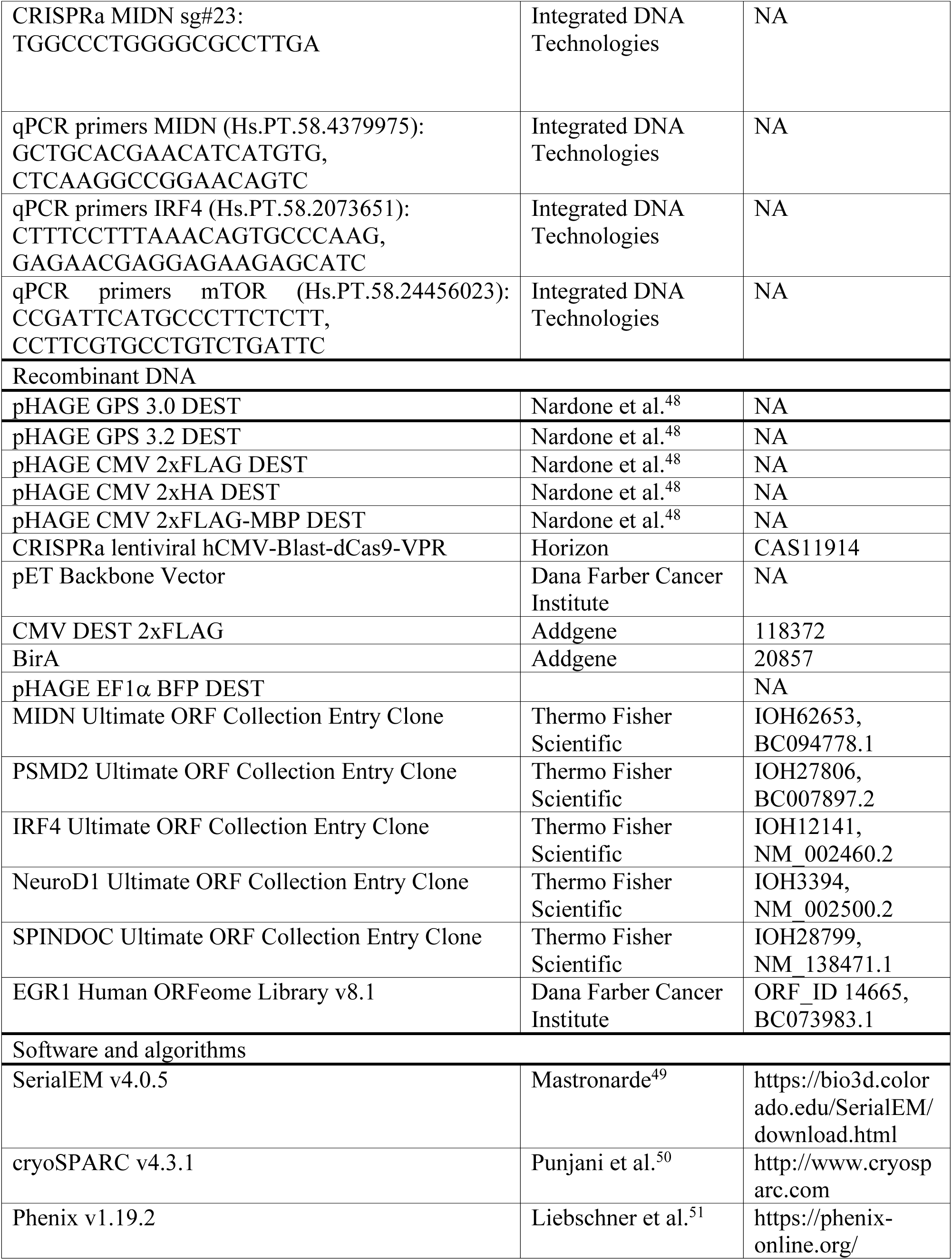
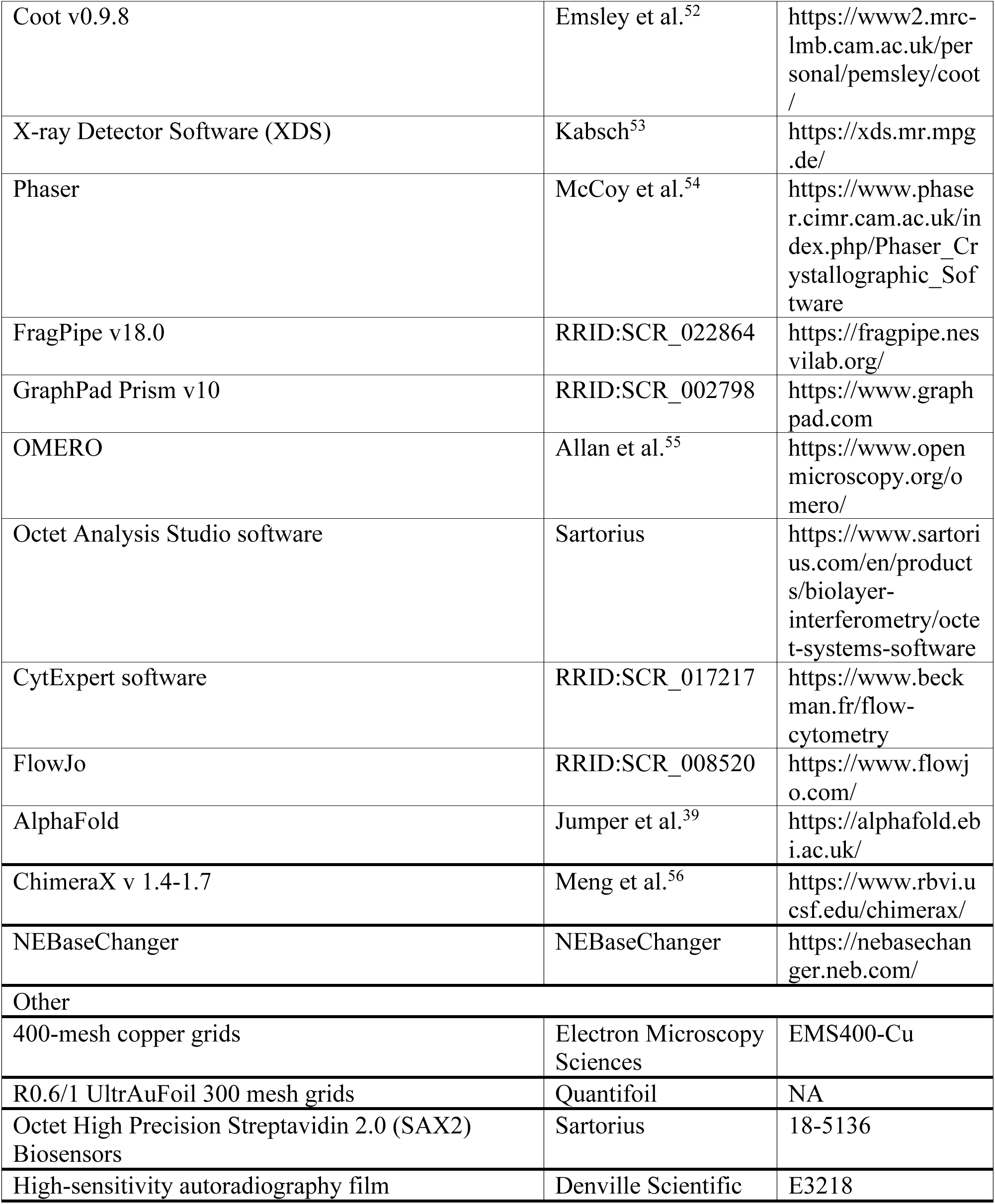

### Resource availability

#### Lead contact

Further information and requests for reagents should be directed to and will be fulfilled by the lead contact, Xin Gu.

#### Data and materials availability

EM maps and models are available under accession numbers PDB 9BUI, PDB 9BV1, PDB 9BV2, PDN 9BV3, EMD-44910, EMD-44927, EMD-44928, EMD-44929, and EMD-44930. The X-ray model is available under accession number PDB 8VQP. All other data are available in the main text or supplemental information.

#### Experimental model and subject details

Human embryonic kidney (HEK)-293T (ATCC, CRL-3216, RRID: CVCL_0063) were cultured in Dulbecco’s Modified Eagle’s Medium (DMEM) (Thermo Fisher Scientific, 11965118) containing 100 units/mL of penicillin and 0.1 mg/mL of streptomycin (Thermo Fisher Scientific, 15070063) and 10% fetal bovine serum (Cytiva, SH30088.03).

AMO-1 (DSMZ, ACC 538), JJN3 (DSMZ, ACC 541), KMS27 (JCRB, JCRB1188), KMS26 (JCRB, JCRB1187), ANBL6 (Sigma, SCC429), MOLP8 (DSMZ, ACC 569), H929 (ATCC, CRL-3580), BSWM1 (gift from Steven Treon’s Lab at DFCI), RAMOS (ATCC, CRL-1596), MWCL1 (gift from Stephen M. Ansell’s Lab at Mayo Clinic), SU-DHL-8 (ATCC, CRL-2961), and Toledo (ATCC, CRL-2631) cells were cultured in Roswell Park Memorial Institute (RPMI) 1640 medium (Thermo Fischer Scientific, A1049101) containing 100 units/mL of penicillin and 0.1 mg/mL of streptomycin, 10% fetal bovine serum, and Plasmocin (InvivoGen, ant-mpp). Cells were kept at 37°C and 5% CO_2_. In the indicated experiments, cells were treated with 10 µM MG132 for 4 hours from 1000x stock solution in dimethyl sulfoxide (DMSO).

### Method details

#### Plasmids

As previously described^5^, plasmid cDNAs containing a stop codon and conferring kanamycin resistance were obtained from the human Ultimate ORF collection (Thermo Fisher Scientific) in the form of a Gateway entry clone: MIDN (IOH62653, BC094778.1), PSMD2 (IOH27806, BC007897.2) IRF4 (IOH12141, NM_002460.2), NeuroD1 (IOH3394, NM_002500.2), and SPINDOC (IOH28799, NM_138471.1). The EGR1 Gateway entry clone (ORF_ID no. 14665, BC073983.1) lacking a stop codon and conferring spectinomycin resistance was obtained from the Human ORFeome library V8.1 (Dana Farber Cancer Institute). The entry clone for *D. gyrociliatus* midnolin was generated by synthesis.

The entry clone for the αHelixC (residues 360 to 432) used in sufficiency experiments was generated by polymerase chain reaction (PCR) using primers containing the following overhangs for cloning into pDONR221 via a BP reaction (Thermo Fisher Scientific, 11789020): attB1, GGGGACAAGTTTGTACAAAAAAGCAGGCTTAgccacc; attB2, GGGGACCACTTTGTACAAGAAAGCTGGGTA. EGR1 residues 113 to 172 were N-terminally appended to midnolin by PCR to generate an entry clone.

Mutations to entry clones were introduced by polymerase chain reaction (PCR) using the Q5 Site-Directed Mutagenesis kit (NEB, E0554S). The PCR primers were designed using the NEBaseChanger online tool. Wild type and mutant versions of entry clones were subcloned into the following destination vectors via an LR reaction (Thermo Fisher Scientific, 11791100): pHAGE CMV 2xFLAG-destination vector for N-terminally tagging MIDN, IRF4, NeuroD1, and SPINDOC, a pHAGE CMV 2xHA-destination vector for N-terminally tagging PSMD2, a CMV-C-2xFLAG destination vector (Addgene, 118372) for C-terminally tagging EGR1, a pHAGE CMV 2xFLAG-MBP-destination vector for N-terminally tagging the midnolin α-HelixC, a GPS 3.0 destination vector for GFP-IRF4, a GPS 3.2 destination vector for EGR1-GFP, or a pHAGE EF1α-destination vector (blue fluorescent protein, BFP) for expressing untagged midnolin for flow cytometry experiments.

The CRISPRmod CRISPRa lentiviral hCMV-Blast-dCas9-VPR plasmid was purchased (Horizon, CAS11914). Below is the information for the sgRNAs expressed using a U6 promoter for the CRISPR-activation experiments:

Non-targeting control: TGTCGTGATGCGTAGACGG

MIDN sg#11: GTGCCGCGGCGGCTCTCAT

MIDN sg#23: TGGCCCTGGGGCGCCTTGA

FLAG-MsyB-αHelixC (WT, L395D, and R405/406/409/410/413D), TNPO1, KPNA2, and the EGR1-Catch domain oligonucleotides codon-optimized for expression in bacteria were generated by synthesis from Integrated DNA Technologies (IDT) to contain overhangs for Gibson assembly (NEB, E2611S). A pET 6xHis-3C bacterial expression vector obtained from the Dana-Farber Cancer Institute (DFCI) Crystallography core facility was digested with BamHI and NotI for direct Gibson assembly of the synthetic oligonucleotides. FLAG-PSMD2 was cloned into a pRK5 vector for expression and purification from bacteria.

2xFLAG-EGR1-midnolin (for cryo-EM):

MDYKDDDDKDYKDDDDKGGGGSGGGGFETSLYKKAGLATMEKVLVETSYPSQTTRLPPITYTGRFSLEPAPNSGNTLWPEPLFSLVSGLVSMTNPPASSSEPQPGGARSCRRGAPGGACELGPAAEAAPMSLAIHSTTGTRYDLAVPPDETVEGLRKRLSQRLKVPKERLALLHKDTRLSSGKLQEFGVGDGSKLTLVPTVEAGLMSQASRPEQSVMQALESLTETQVSDFLSGRSPLTLALRVGDHMMFVQLQLAAQHAPLQHRHVLAAAAAAAAARGDPSIASPVSSPCRPVSSAARVPPVPTSPSPASPSPITAGSFRSHAASTTCPEQMDCSPTASSSASPGASTTSTPGASPAPRSRKPGAVIESFVNHAPGVFSGTFSGTLHPNCQDSSGRPRRDIGTILQILNDLLSATRHYQGMPPSLAQLRCHAQCSPASPAPDLAPRTTSCEKLTAAPSASLLQGQSQIRMCKPPGDRLRQTENRATRCKVERLQLLLQQKRLRRKARRDARGPYHWSPSRKAGRSDSSSSGGGGSPSEASGLGLDFEDSVWKPEANPDIKSEFVVA

D. gyrociliatus midnolin:

MNVNLIICPSTGGEYRLSVDLGENVNDLRRSVASKFRLDAERIRLIYNDRNLTTGTLEENDVGENSKIILLPCLRSGLVPKSVESTSVQIAEALKTLSGKQIDDFLCGRKPLNLRLILGGHFVYVQLQLSSTSKCPKQPSRIEDVRCHGSGVFSGTFSGTVRSPKSKNGASLVAGILNDILFSQRRRRDDQMTGEIRRAASRHSSPDKQRRVEDDQTRSKMAALKEALVERRRKKRKFSPY DRLLV

The following proteins were expressed in bacteria using a pET plasmid:

FLAG-MsyB-αHelixC (wild type):

MQLSHHHHHHSSGLEVLFQGPGGGSDYKDDDDKGGGGSMYATLEEAIDAAREEFLADNPGIDAEDANVQQFNAQKYVLQDGDIMWQVEFFADEGEEGECLPMLSGEAAQSVFDG DYDEIEIRQEWQEENTLHEWDEGEFQLEPPLDTEEGRAAADEWDERETSLYKKAGLATSASLLQGQSQIRMCKPPGDRLRQTENRATRCKVERLQLLLQQKRLRRKARRDARGPYHWSPSRKAGRSDSSSS

FLAG-MsyB-αHelixC (L395D):

MQLSHHHHHHSSGLEVLFQGPGGGSDYKDDDDKGGGGSMYATLEEAIDAAREEFLADNPGIDAEDANVQQFNAQKYVLQDGDIMWQVEFFADEGEEGECLPMLSGEAAQSVFDGDYDEIEIRQEWQEENTLHEWDEGEFQLEPPLDTEEGRAAADEWDERETSLYKKAGLATSASLLQGQSQIRMCKPPGDRLRQTENRATRCKVERDQLLLQQKRLRRKARRDARGPYHWSPSRKAGRSDSSSS

FLAG-MsyB-αHelixC (5RD):

MQLSHHHHHHSSGLEVLFQGPGGGSDYKDDDDKGGGGSMYATLEEAIDAAREEFLADNPGIDAEDANVQQFNAQKYVLQDGDIMWQVEFFADEGEEGECLPMLSGEAAQSVFDGDYDEIEIRQEWQEENTLHEWDEGEFQLEPPLDTEEGRAAADEWDERETSLYKKAGLATSASLLQGQSQIRMCKPPGDRLRQTENRATRCKVERLQLLLQQKRLDDKADDDADGPYHWSPSRKAGRSDSSSS

FLAG-Avi-MsyB-αHelixC (WT):

MQLSHHHHHHSSGLEVLFQGPGGGSDYKDDDDKGGGGGLNDIFEAQKIEWHESSGGSMYATLEEAIDAAREEFLADNPGIDAEDANVQQFNAQKYVLQDGDIMWQVEFFADEGEEGECLPMLSGEAAQSVFDGDYDEIEIRQEWQEENTLHEWDEGEFQLEPPLDTEEGRAAADEWDERETSLYKKAGLATSASLLQGQSQIRMCKPPGDRLRQTENRATRCKVERLQLLLQQKRLRRKARRDARGPYHWSPSRKAGRSDSSSS

FLAG-Avi-MsyB-αHelixC (5RD):

MQLSHHHHHHSSGLEVLFQGPGGGSDYKDDDDKGGGGGLNDIFEAQKIEWHESSGGSMYATLEEAIDAAREEFLADNPGIDAEDANVQQFNAQKYVLQDGDIMWQVEFFADEGEE GECLPMLSGEAAQSVFDGDYDEIEIRQEWQEENTLHEWDEGEFQLEPPLDTEEGRAAADEWDERETSLYKKAGLATSASLLQGQSQIRMCKPPGDRLRQTENRATRCKVERLQLLLQ QKRLDDKADDDADGPYHWSPSRKAGRSDSSSS

TNPO1:

MQLSHHHHHHSSGLEVLFQGPGGGVWDRQTKMEYEWKPDEQGLQQILQLLKESQSPDTTIQRTVQQKLEQLNQYPDFNNYLIFVLTKLKSEDEPTRSLSGLILKNNVKAHFQNFPNGVTDFIKSECLNNIGDSSPLIRATVGILITTIASKGELQNWPDLLPKLCSLLDSEDYNTCEGAFGALQKICEDSAEILDSDVLDRPLNIMIPKFLQFFKHSSPKIRSHAVACVNQFIISRTQALMLHIDSFIENLFALAGDEEPEVRKNVCRALVMLLEVRMDRLLPHMHNIVEYMLQRTQDQDENVALEACEFWLTLAEQPICKDVLVRHLPKLIPVLVNGMKYSDIDIILLKGDVEEDETIPDSEQDIRPRFHRSRTVAQQHDEDGIEEEDDDDDEIDDDDTISDWNLRKCSAAALDVLANVYRDELLPHILPLLKELLFHHEWVVKESGILVLGAIAEGCMQGMIPYLPELIPHLIQCLSDKKALVRSITCWTLSRYAHWVVSQPPDTYLKPLMTELLKRILDSNKRVQEAACSAFATLEEEACTELVPYLAYILDTLVFAFSKYQHKNLLILYDAIGTLADSVGHHLNKPEYIQMLMPPLIQKWNMLKDEDKDLFPLLECLSSVATALQSGFLPYCEPVYQRCVNLVQKTLAQAMLNNAQPDQYEAPDKDFMIVALDLLSGLAEGLGGNIEQLVARSNILTLMYQCMQDKMPEVRQSSFALLGDLTKACFQHVKPCIADFMPILGTNLNPEFISVCNNATWAIGEISIQMGIEMQPYIPMVLHQLVEIINRPNTPKTLLENTAITIGRLGYVCPQEVAPMLQQFIRPWCTSLRNIRDNEEKDSAFRGICTMISVNPSGVIQDFIFFCDAVASWINPKDDLRDMFCKILHGFKNQVGDENWRRFSDQFPLPLKERLAAFYGV

KPNA2:

MQLSHHHHHHSSGLEVLFQGPGGGGLNDIFEAQKIEWHESSGLEVLFQGPGSSTNENANTPAARLHRFKNKGKDSTEMRRRRIEVNVELRKAKKDDQMLKRRNVSSFPDDATSPLQENRNNQGTVNWSVDDIVKGINSSNVENQLQATQAARKLLSREKQPPIDNIIRAGLIPKFVSFLGRTDCSPIQFESAWALTNIASGTSEQTKAVVDGGAIPAFISLLASPHAHISEQAVWALGNIAGDGSVFRDLVIKYGAVDPLLALLAVPDMSSLACGYLRNLTWTLSNLCRNKNPAPPIDAVEQILPTLVRLLHHDDPEVLADTCWAISYLTDGPNERIGMVVKTGVVPQLVKLLGASELPIVTPALRAIGNIVTGTDEQTQVVIDAGALAVFPSLLTNPKTNIQKEATWTMSNITAGRQDQIQQVVNHGLVPFLVSVLSKADFKTQKEAVWAVTNYTSGGTVEQIVYLVHCGIIEPLMNLLTAKDTKIILVILDAISNIFQAAEKLGETEKLSIMIEECGGLDKIEALQNHENESVYKASLSLIEKYFSVEEEEDQNVVPETTSEGYTFQVQDGAPGTFNF

EGR1-Catch domain (for crystallography):

MQLSHHHHHHSSGLEVLFQGPGGSPPITYTGRFSLEPAPNSGNTLSRPEQSVMQALESLTETQVSDFLSGRSPLTLALRVGDHMMFVQLQLAAQHAPPAPRSRKPGAVIESFVNHAPGVFSGTFSGTLHPNCQDSSGRPRRDIGTILQILNDLLSATRHYQGMPPSLAQLRCHA

PSMD2-Avi-FLAG:

MEEGGRDKAPVQPQQSPAAAPGGTDEKPSGKERRDAGDKDKEQELSEEDKQLQDELEMLVERLGEKDTSLYRPALEELRRQIRSSTTSMTSVPKPLKFLRPHYGKLKEIYENMAPGE NKRFAADIISVLAMTMSGERECLKYRLVGSQEELASWGHEYVRHLAGEVAKEWQELDDAEKVQREPLLTLVKEIVPYNMAHNAEHEACDLLMEIEQVDMLEKDIDENAYAKVCLYLT SCVNYVPEPENSALLRCALGVFRKFSRFPEALRLALMLNDMELVEDIFTSCKDVVVQKQMAFMLGRHGVFLELSEDVEEYEDLTEIMSNVQLNSNFLALARELDIMEPKVPDDIYKTHLENNRFGGSGSQVDSARMNLASSFVNGFVNAAFGQDKLLTDDGNKWLYKNKDHGMLSAAASLGMILLWDVDGGLTQIDKYLYSSEDYIKSGALLACGIVNSGVRNECDPALALLSDYVLHNSNTMRLGSIFGLGLAYAGSNREDVLTLLLPVMGDSKSSMEVAGVTALACGMIAVGSCNGDVTSTILQTIMEKSETELKDTYARWLPLGLGLNHLGKGEAIEAILAALEVVSEPFRSFANTLVDVCAYAGSGNVLKVQQLLHICSEHFDSKEKEEDKDKKEKKDKDKKEAPADMGAHQGVAVLGIALIAMGEEIGAEMALRTFGHLLRYGEPTLRRAVPLALALISVSNPRLNILDTLSKFSHDADPEVSYYSIFAMGMVGSGTNNARLAAMLRQLAQYHAKDPNNLFMVRLAQGLTHLGKGTLTLCPYHSDRQLMSQVAVAGLLTVLVSFLDVRNIILGKSHYVLYGLVAAMQPRMLVTFDEELRPLPVSVRVGQAVDVVGQAGKPKTITGFQTHTTPVLLAHGERAELATEEFLPVTPILEGFVILRKNPNYDLGGGGSGLNDIFEAQKIEWHEGGGGSDYKDDDDK

#### Purification of the Midnolin-Proteasome Complex

Four purifications were performed for cryo-EM analysis. (1) The lysis buffer lacked ATP and magnesium, and the bait was wild-type 2xFLAG-midnolin. (2) The lysis buffer contained both ATP and magnesium, and the bait was wild-type 2xFLAG-midnolin. (3) The lysis buffer contained both ATP and magnesium, and the bait was G83E 2xFLAG-midnolin. (4) The lysis buffer contained both ATP and magnesium, and the bait was 2xFLAG-EGR1-midnolin.

3.5 million HEK-293T cells were seeded into 50 15 cm plates per prep. Two days after seeding, the cells were transfected using PolyJet with 10 µg DNA per plate of 2xFLAG-midnolin under the control of a CMV promoter (500 µL DMEM + DNA and 500 µL DMEM + 30 µL PolyJet per plate). The media was changed one day after the transfection. Two days post-transfection, the cells were rinsed with ice-cold PBS and harvested by scraping in 1 mL of lysis buffer containing 40 mM HEPES, 100 mM NaCl, 2 mM MgCl_2_, 5 mM ATP, 1x HALT protease, 0.5% CHAPS per 15 cm plate. The lysate was incubated end-to-end for 20 minutes at 4°C before centrifuging at 21,000 x g. The clarified lysate was poured into two 50 mL falcon tubes. Using the lysis buffer, 1.8 mL of packed M2 FLAG agarose resin (Millipore, A2220) was rinsed four times in 2 mL Eppendorf tubes. Note that the FLAG M2 resin may change to a yellow/light brown color because of the HALT cocktail. The washed resin was added to the 50 mL falcons containing the clarified lysate and rotated end-to-end for 1.5 hours at 4°C. The falcon lids were sealed with parafilm to ensure there was no leaking. After incubating, the resin-containing cell lysate was flowed through a column (BioRad, 7321010) at 4°C by gravity. The two 50 mL falcons were rinsed with 15 mL lysis buffer for each falcon to wash off residual FLAG resin and the solution was flowed through the column. Finally, the resin on the column was rinsed once with 20 mL of lysis buffer, followed by 20 mL of size-exclusion chromatography (SEC) buffer containing 40 mM HEPES, 100 mM NaCl, 2mM MgCl_2_, 5mM ATP, 0.05% CHAPS. The proteins were eluted from the resin using SEC buffer supplemented with 0.5 mg/mL of 3xFLAG peptide (APExBIO, A6001). Specifically, the resin on the column was incubated with 1.8 mL of elution buffer for 20 minutes at room temperature and the eluate was collected in a fresh tube. This process was repeated for a total of four elutions. To determine if protein was present in the elutions, an absorbance measurement at 280 nm was recorded using the elution buffer as a blank to determine if protein was present in the elutions.

The elutions containing proteins were pooled and concentrated (3,500 x g) using an Amicon Ultra-15, 10 kDa MWCO concentrator until ∼500 µL. It was important to pre-wet the filter using the SEC buffer before adding protein samples and to pipet up and down gently between centrifuging rounds to prevent unnecessary crashing. The concentration before SEC was around 4-5 mg/mL. The protein was centrifuged at 21,000 x g for 1 minute at 4°C to remove any debris. A Superose6 column was equilibrated using the SEC buffer and a 0.5 mL injection loop was washed thoroughly with the SEC buffer. The loop was then manually injected with 0.5 mL of concentrated eluted protein. The 0.5 mL fractions corresponding to the protein of interest were collected, pooled, and concentrated using Amicon Ultra-15, 10 kDa MWCO (4,800 x g, 6 minutes intervals) to a final volume of around 300 µL at 3-4 mg/mL of pure protein. The purified proteins were used directly for negative stain electron microscopy and cryo-EM sample preparation to prevent freezing and thawing of the sample.

#### Negative Stain Electron Microscopy

3.5 µl of midnolin-proteasome complexes purified as above was applied to glow-discharged (30 seconds, 30 mA) 400-mesh copper grids (EMS400-Cu) coated with a continuous layer of approximately 10 nm carbon. After 1 minute, the grid was blotted using filter paper (VMR, 28310-081), immediately stained with 1.5 % uranyl formate solution (Electron Microscopy Science, 22451), and blotted again. The staining procedure was repeated twice with a 1-minute incubation with uranyl formate before the final blotting step. The grid was then air-dried at room temperature before imaging. Images were collected using a FEI Tecnai T12 transmission electron microscope operating at 120 kV and equipped with a Gatan UltraScan 895 (4k x 4k) CCD camera at a nominal magnification of 67,000× corresponding to a calibrated pixel size of 1.68 Å and a defocus value of -3.0 µm.

#### Cryo-EM Sample Preparation and Data Collection

3.5 µl of 4 mg/mL midnolin-proteasome complexes purified as above was applied to glow-discharged R0.6/1 UltrAuFoil 300 mesh grids (Quantifoil) and frozen in liquid ethane using a Vitrobot Mark IV (Thermo Fisher Scientific) set at 4°C and 100% humidity with a 10 second wait time, 3 second blot time, and +8 blot force. All datasets were collected using a Titan Krios (Thermo Fisher Scientific) operating at 300 kV and equipped with a BioQuantum K3 imaging filter with a 20-eV slit width and a K3 summit direct electron detector (Gatan) in counting mode at a nominal magnification of 105,000× corresponding to a calibrated pixel size of 0.825 Å. Semi-automated data collection was performed with SerialEM v4.0.5. For the WT MIDN -ATP dataset, a 2.597-second exposure was fractionated into 50 frames, resulting in a total exposure of 52.98 e-/Å^2^. The defocus targets were −1.0 to −2.2 µm. For the WT MIDN +ATP dataset, a 2.652-second exposure was fractionated into 48 frames, resulting in a total exposure of 50.54 e-/Å^2^. The defocus targets were −1.0 to −2.2 µm. For the G83E MIDN +ATP dataset, a 2.65-second exposure was fractionated into 50 frames, resulting in a total exposure of 53 e-/Å^2^. The defocus targets were −1.0 to −2.2 µm. For the EGR1-MIDN +ATP dataset, a 3.02-second exposure was fractionated into 54 frames, resulting in a total exposure of 53.79 e-/Å^2^. The defocus targets were −0.8 to −1.8 µm.

#### Image Processing and Model Building

Data processing for all 4 datasets was performed in cryoSPARC v4.3.1 (**Figure S2**). After patch-based motion correction and CTF estimation, micrographs with severe contamination or poor CTF fits were removed. 3,914 (prep 1, WT MIDN -ATP), 3,396 (prep 2, WT MIDN +ATP), 4,095 (prep 3, G83E MIDN +ATP) or 8,035 (prep 4, EGR1-MIDN +ATP) micrographs were subjected to automated particle picking using templates generated from blob-based picking. The particles were extracted with a box size of 800 and downsampled to a box size of 200 for the initial classification steps. After 2D classification, 380,637 (prep 1), 380,842 (prep 2), 285,529 (prep 3), and 658,210 (prep 4) particles were selected for heterogeneous refinement using multiple reference volumes generated by ab initio reconstruction. Particles in the best classes were subjected to homogeneous refinement with C2 symmetry, followed by symmetry expansion. Afterward, all particles were re-centered on the 19S particle, extracted with a box size of 440, and downsampled to a box size of 220. Good 19S particles after 2D classification were then subjected to multiple rounds of heterogeneous refinements.

35,838 particles from prep 1, 2, and 3 datasets in classes with density corresponding to the Ubl domain were unbinned, local motion corrected, combined, and subjected to non-uniform and local refinement. 340,023 particles from the EGR1-MIDN +ATP dataset with a similar AAA+ ATPase state but no obvious Ubl density were subjected to 3D classification with a mask focused on PSMD2. 113,946 particles in the class with the clearest PSMD2 density were unbinned, local motion corrected and subjected to non-uniform and local refinement. 228,049 particles from the EGR1-MIDN +ATP dataset with clear PSMD2 were subjected to an additional round of heterogeneous refinement to reveal two classes showing the midnolin αHelix-C bound to PSMD2 and different AAA+ ATPase states. The particles from these classes were combined with particles in corresponding classes in prep 1, prep 2, and prep 4 datasets after unbinning, local motion correction, and additional heterogeneous refinement with a mask focused on PSMD2. 186,489 particles in the M2-A state and 87,326 particles in the M2-B state containing clear PSMD2 density were used for final non-uniform and local refinements.

Unsharpened and post-processed maps from cryoSPARC and deepEMhancer were used for interpretation. Local resolution estimation (**Figure S3**) and 3D variability analysis were performed in cryoSPARC. An initial model for the M2-A conformation was generated primarily by rigid body fitting individual AlphaFold models of each proteasomal subunit, the AlphaFold multimer model of PSMD2 with the midnolin αhelix-C, cross-referencing the output from ModelAngelo^57^, particularly for the AAA+ ATPase subunits. The model was manually adjusted in *Coot* v.0.9.8 in between iterative rounds of real space refinement in Phenix v.1.19.2. The M2-A model was then used as an initial model for the M2-B conformation, which was refined the same way after adjustment of the AAA+ ATPase subunit conformations in *Coot* guided by the output from ModelAngelo. The M1 conformations were modeled in the same way, starting with rigid body fitting of individual AlphaFold models, including the AlphaFold-multimer model of PSMD14 with the midnolin Ubl domain, and cross-referencing with the output from ModelAngelo. Identification and placement of ZFAND5 in the M1 w/Ubl conformation was guided by PDB 7QXW^23^. Figures were made using ChimeraX v.1.5 – v.1.7.

#### Crystallography

##### Protein expression and purification

The chimeric EGR1-Catch domain construct was overexpressed in E. coli BL21 (DE3) and purified using affinity chromatography and size-exclusion chromatography. Briefly, cells were grown at 37°C in TB medium in the presence of 50 μg/mL of kanamycin to an OD of 0.8, cooled to 17°C, induced with 400 μM isopropyl-1-thio-D-galactopyranoside (IPTG), incubated overnight for 20 hours at 17°C, collected by centrifugation, and stored at -80°C. Cell pellets were lysed in buffer A (25 mM HEPES, pH 7.5, 500 mM NaCl, and 1 mM TCEP) using a Microfluidizer (Microfluidics), and the resulting lysate was centrifuged at 30,000 x g for 40 minutes. Ni-NTA beads (Qiagen) were mixed with cleared lysate for 45 minutes and washed with buffer A. Beads were transferred to an FPLC-compatible column, and the bound protein was washed further with buffer A for 10 column volumes and eluted with buffer B (25 mM HEPES, pH 7.5, 500 mM NaCl, 10% glycerol, 0.5 mM TCEP, and 400 mM imidazole). The eluted sample was concentrated and purified further using a Superdex 200 16/600 column (Cytiva) in buffer C containing 20 mM HEPES, pH 7.5, 300 mM NaCl, and 0.5 mM TCEP. 3C protease was added to SEC fractions containing the EGR1-Catch domain, and His-tag was removed by applying it to a second Ni-NTA column. The eluant containing cleaved EGR1-Catch domain protein was concentrated to ∼15 mg/mL and stored at -80°C.

##### Crystallization

A solution of 15 mg/mL EGR1-Catch domain was crystallized in 1.4 M ammonium sulfate and 100 mM HEPES, pH 7.0 by sitting-drop vapor diffusion at 27°C. Crystals were transferred briefly into a crystallization buffer containing 25% glycerol before flash-freezing in liquid nitrogen.

##### Determination

Diffraction data were collected at beamline 17-ID-2 of the National Synchrotron Light Source II (Brookhaven National Laboratory). Datasets were integrated and scaled using X-ray Detector Software (XDS)^53^. Structures were solved by molecular replacement using the program Phaser^54^ and the AlphaFold-predicted model as a search model. Iterative manual model building and refinement using Phenix^51^ and *Coot*^52^ led to a model with excellent statistics.

#### Purification of Importins and MsyB-tagged αhelix-C

20 µL of chloramphenicol-resistant Rosetta (DE3) cells (Millipore, 71397-3) were transformed with pET plasmid conferring kanamycin resistance and IPTG-inducible expression of 6xHis-FLAG-MsyB-αhelix-C, -TNPO1, or -KPNA2. For *in vivo* biotinylation of FLAG-Avi-MsyB-αhelix-C, cells were co-transformed with a modified BirA plasmid (Addgene, 20857) lacking the His-tag. Transformed cells were plated onto chloramphenicol and kanamycin +/- ampicillin and grown at 37°C overnight. The next day, single colonies were picked into 50 mL LB starter cultures containing the appropriate antibiotic, and the bacteria were grown overnight at 37°C with 200 rpm shaking. The next day, the starter cultures were transferred to 1 L of fresh LB containing the appropriate antibiotic, and the bacteria were grown at 37°C to an O.D. = 0.6. Once reaching this density, the bacteria were cooled at 4°C for 10 minutes, and 0.5 mM isopropyl β- d-1- thiogalactopyranoside IPTG (Sigma, I5502-10G) was added directly to the liquid cultures. For biotinylation, the bacteria were also supplemented with 50 µM biotin. The bacteria were then grown at 16°C for 16-18 hours to induce protein expression.

Post induction, the bacteria were transferred to 1 L centrifuge bottles and pelleted at 4,000 x g for 15 minutes at 4°C. The pellet was resuspended in 35 mL of buffer containing 50 mM HEPES pH 7.4, 150 mM NaCl, and 10 mM imidazole (Sigma, 68268-500ML-F). The resuspended bacteria were again pelleted at 4,000 x g for 15 minutes at 4°C. The supernatant was removed, and the cells were resuspended in 35 mL of lysis buffer containing 50 mM HEPES pH 7.4, 150 mM NaCl, 10 mM imidazole, 1 mM dithiothreitol DTT (Thermo Fisher Scientific, R0862), benzonase (Millipore, 71206-3), and protease inhibitor tablets (Sigma, 11873580001). The cells were lysed using a microfluidizer (700 bar, 4 cycles, 20 mL/minute flow rate). Lysate was clarified by centrifuging at 21,000 x g for 20 minutes at 4°C. While centrifuging, 0.8 mL column volume (1.6 mL total volume per protein prep) of Ni-NTA agarose (Qiagen, 30210) was washed three times with lysis buffer. The supernatant was then incubated with the washed Ni-NTA agarose in a 50 mL conical tube for 30 minutes with end-to-end rotation at 4°C. After incubation, the mixture was subjected to gravity flow and both the conical tube, and the columns were rinsed with 50-60 mL total buffer at 4°C. The protein was eluted 4 times from the resin by sequentially incubating for 5 minutes with 500 µL of buffer containing 50 mM HEPES, 150 mM NaCl, and 200 mM imidazole. The absorbance A280 of each elution was measured using a nanodrop.

The elutions containing the highest amount of protein, usually elutions 1, 2, and 3, were pooled and concentrated to 500 µL using Amicon Ultra-15, 3 kDa MWCO (Millipore, UFC9003). The protein was centrifuged at 21,000 x g for 1 minute to remove any debris. Superdex 200 size-exclusion chromatography was equilibrated using a buffer containing 50 mM HEPES, 150 mM NaCl, and 1 mM DTT. A 0.5 mL injection loop was washed thoroughly with this same buffer and the loop was manually injected with 0.5 mL of concentrated eluted protein. The 0.5 mL fractions corresponding to the protein of interest were collected, pooled, and concentrated using Amicon Ultra-15, 3 kDa MWCO to a final volume of around 300-400 µL at 3-8 mg/mL of pure protein.

#### Purification of Human Proteasome

HBTH-Rpn11 suspension HEK-293 cells^58^ were grown in 600 mL of Expi293 expression media (Thermo Fischer Scientific, A1435101) to a final concentration of 5 million cells/mL. Cells were harvested by spinning at 1000 x g for 5 minutes. The media was removed, and the cell pellet was rinsed once with 100 mL of PBS before centrifuging again at 1000 x g for 5 minutes. The PBS was removed, and the cells were resuspended in 70 mL of lysis buffer containing 40 mM HEPES pH 7.4, 100 mM NaCl, 0.5 % CHAPS, 1 mM MgCl_2_, 1 mM DTT, 5 mM ATP, and 1x HALT protease and phosphatase inhibitor. Cells were Dounce homogenized gently for 12 stokes/2 mL of lysis buffer. The homogenate was subsequently incubated at 4°C for 20 minutes with end-to-end rotation. The lysate was then centrifuged at 21,000 x g for 25 minutes at 4°C to remove cell debris. During the spin, prepare ∼2.5 mL slurry of NeutrAvidin Agarose (Thermo Fischer Scientific, 29200) and wash 4 times with lysis buffer to remove the storage buffer. Incubate the resin with the cleared supernatant, rotating end-to-end at 4°C for 2 hours. The beads were rinsed with lysis buffer 3 times, 20 mL each time, with gentle spinning at 2000 x g for 3 minutes at 4°C to precipitate the beads after each wash. The resin was then rinsed twice with 25 mL of TEB buffer containing 50 mM Tris pH 7.5, 50 mM NaCl, 2.5 mM ATP, 1 mM MgCl2, and 1 mM DTT. The beads were then dried with a needle and vacuum before adding 4 mL of TEB containing 18 µL of HisTEV (Thermo Fischer Scientific, 12-575-015). Cleavage was allowed to occur overnight at 4°C with gentle shaking. The eluate (∼4 mL) was collected. The beads were washed with an additional 2.5 mL of TEB buffer. The 6.5 mL pool containing the two elutes was concentrated using an Amicon Ultra-15, 10 kDa MWCO concentrator (Millipore, UFC9010) to a final volume of ∼500 µL containing about 0.6 mg/mL of human proteasomes. The solution was aliquoted and snap-frozen in liquid nitrogen for storage at -80°C.

#### Coomassie Staining

Tris-Glycine gels soaked in Coomassie Brilliant Blue R-250 staining solution (BioRad, 1610436) were microwaved for 15 seconds and incubated at room temperature with gentle rocking for 30 minutes. Gels were then destained using 35 % methanol and 10 % acetic acid in water in the presence of a Kimwipe with gentle rocking overnight.

#### Lentivirus Production

HEK-293T cells were seeded into 6-well culture plates at 300,000 cells/well. The cells were allowed to grow for 2 days before they were transfected using PolyJet (SignaGen, SL100688) following the manufacturer’s instructions. Briefly, plasmids encoding Tat, Rev, Gag-Pol, and VSV-G were mixed equimolar with lentiviral transfer vectors. Specifically, 1 µg of total plasmid DNA was diluted into DMEM lacking supplements. Separately, 3 µL of PolyJet reagent was diluted into DMEM lacking supplements. The PolyJet solution was added to the DNA solution. The mixture was incubated for 15 minutes at room temperature and added to cells dropwise. The media was replaced with fresh media one day post-transfection. The lentiviral supernatant was collected at 48 hours post-transfection, passed through a 0.45 µm filter, and applied directly onto cells.

#### CRISPR-activation Cell-Titer-Glow Assay

AMO-1 cells stably expressing nuclease-deactivated Cas9-VPR were transduced by incubating 1.5 million cells/well in a 12-well plate with lentivirus (sgRNA + RFP) and Polybrene and spinfecting at 800 x g for 90 minutes. The media was carefully removed after spinfection and replaced with fresh media. The following day, cells were expanded in 5 mL of fresh media in a T25 flask. Three days post-spinfection, the cells were analyzed by flow cytometry for the percentage of RFP+. We used the minimal dose of lentivirus to achieve 100% RFP+ cells. Before attempting this experiment, lentiviral particles should be titered to determine the minimal dose needed for 100% positive cells. The cells were then diluted to 50,000 cells/mL, and 100 µL (5,000 cells) was aliquoted into opaque 96-well plates, one plate for each day of measuring proliferation. On the day of seeding (day 0), 100 µL of Cell Titer Glow reagent (Promega, G9242) was added to each well using a multichannel and incubated for 5 minutes at room temperature. Luminescence was measured using a plate reader. Proliferation measurements in replicate 96-well plates were obtained every 2 days for 8 days total.

#### Immunoprecipitation

HEK-293T cells were seeded at 1 million cells per 10 cm culture dish. Two days post-seeding, the cells were transfected with 3 µg of the indicated plasmid using PolyJet. The media was replaced with fresh media one day post-transfection. Two days post-transfection, the cells were rinsed once with ice-cold PBS and 700 µL of lysis buffer containing 0.5 % CHAPS, 100 mM NaCl, 40 mM HEPES pH 7.4, 1x protease-phosphatase inhibitor cocktail (Thermo Fisher Scientific, 78441) was applied directly to the dish. The cells were harvested into Eppendorf tubes by scraping and lysates were incubated with end-to-end rotation for 15-20 minutes at 4°C. The tubes were then centrifuged at 21,000 x g for 15 minutes at 4°C. During the centrifuge step, 15 µL/plate of anti-FLAG (Sigma, M8823, RRID: AB_2637089) or anti-HA (Thermo Fisher Scientific, 88836, RRID: AB_2749815) magnetic beads were rinsed 3 times using 700 µL of lysis buffer. After centrifuging, a 30 µL aliquot of whole-cell lysate was collected into 200 µL of Tris-Glycine SDS sample buffer (Thermo Fisher Scientific, LC2676) containing 10% 2-mercaptoethanol. The remainder of the supernatant was incubated with the magnetic beads end-to-end for 1 hour at 4°C. The beads were then rinsed 3 times using 700 µL of lysis buffer and resuspended in 50 µL of Tris-Glycine SDS sample buffer. The protein samples were then denatured for 3 minutes at 95°C. The IP samples were diluted 1:20 in Tris-Glycine SDS sample buffer to blot for the bait protein. 15 µL of whole-cell lysate or IP sample was loaded to blot for co-immunoprecipitated proteins.

#### Immunoblotting

Protein samples were resuspended in Tris-Glycine SDS sample buffer (Thermo Fisher Scientific, LC2676) containing 10% 2-mercaptoethanol and denatured for 3 minutes at 95°C. The samples were then loaded into 4-12% Tris-Glycine 15-well pre-cast gels (Thermo Fisher Scientific, XP04125BOX) and electrophoresis was carried out at a constant 165 volts in 1x Tris-Glycine SDS running buffer (Thermo Fisher Scientific, LC2675-4) until the molecular weight ladder (Thermo Fisher Scientific, 26619) reached the bottom of the gel. Proteins in the gel were transferred to a 0.2 µm nitrocellulose membrane (BioRad, 170-4158) using the Trans-Blot Turbo Transfer System (BioRad). The protein-containing membranes were blocked using 5% milk (LabScientific, M-0842) in 1x TBST (CST, 9997S) for 15-30 minutes with gentle rocking at room temperature. All primary antibodies were diluted directly in the 5% milk blocking solution at a 1:1000 dilution by volume and incubated overnight with gentle rocking at 4°C.

The following primary antibodies were used: rabbit anti-PSMD2 (CST, 25430, RRID: AB_2798903), rabbit anti-PSMA2 (CST, 2455, RRID: AB_2171400), rabbit anti-PSMB5 (CST, 12919, RRID: AB_2798061), rabbit anti-HA (CST, 3724, RRID: AB_1549585), rabbit anti-FLAG (CST, 14793, RRID: AB_2572291), rabbit anti-mTOR (CST, 2983, RRID: AB_2105622), rabbit anti-TNPO1 (CST, 31452, RRID: AB_3662993), rabbit anti-KPNA2 (CST, 14372, RRID: AB_2798467), rabbit anti-IRF4 (CST, 4299, RRID: AB_10547141).

After incubating overnight in primary antibody, the membranes were rinsed 3 times using 1x TBST as quick washes. The membranes were also incubated in 1x TBST for three 10-minute washes with gentle rocking. After these rinses, the membranes were incubated with 5% milk in 1x TBST containing a 1:2000 dilution by volume of anti-rabbit IgG, HRP-linked (CST, 7074, RRID: AB_2099233) secondary antibody. The membranes were incubated in secondary antibody for 1 hour at room temperature with gentle rocking before rinsing as done after the primary antibody incubation. After rinsing, the membranes were exposed to Pierce ECL western blotting substrate (Thermo Fisher Scientific, 32106) for 1-2 minutes at room temperature. High-sensitivity autoradiography film (Denville Scientific, E3218) was used to collect all immunoblotting data.

#### Immunofluorescence

HEK-293T cells were seeded in 6-well culture plates containing Poly-D-Lysine coated coverslips (TED PELLA, Inc.) at 300,000 cells per well. On the following day, the culture media was aspirated, and the cells were rinsed with cold phosphate-buffered saline (PBS) once before fixation using 4% paraformaldehyde (PFA) diluted in PBS for 15 minutes at room temperature. The cells were then rinsed 3 times using PBS and the cells were permeabilized with 0.05% Triton diluted in PBS for 5 minutes at room temperature. The cells were rinsed 3 times with PBS and incubated in immunofluorescence blocking buffer (LI-COR, 927-70001) for 45 minutes at room temperature. The coverslips were then transferred faceup from the 6-well culture plate to a parafilm-coated 15 cm culture dish. Mouse anti-FLAG (Sigma, F1804, RRID: AB_262044) primary antibody was diluted 1:400 in the blocking buffer. The cells facing upward were incubated in 100 µL of the primary antibody solution by applying this solution directly on top of the cells. The cells were incubated in primary antibody overnight at 4°C in the 15 cm culture dish that was kept hydrated with a wet Kimwipe. The following day, the cells were rinsed by gently dipping the coverslips 50 times in and out of a 50 mL conical tube containing PBS. After rinsing, the coverslips were again placed faceup on a parafilm-coated dish. A 100 µL blocking solution containing a 1:500 dilution of anti-mouse Alexa 488-conjugated secondary antibody (Thermo Fisher Scientific, A-11001, RRID: AB_2534069) and 8 µM Hoechst 33342 dye (Thermo Fisher Scientific, H3570) was applied directly on top of the cells. The cells were incubated in this solution for 1 hour at room temperature in the dark. After incubation, the coverslips were again rinsed by gently dipping the coverslips 50 times in and out of a 50 mL conical tube containing PBS. The coverslips were mounted on glass slides using ProLong Glass antifade mountant (Thermo Fisher Scientific, P36982).

Images were acquired using a W1 Yokogawa Spinning disk Nikon Ti inverted confocal microscope with a 50 µm pinhole disk. All images were acquired using a Plan Apo 60X/1.4 oil immersion objective and an Andor Zyla 4.2 Plus sCMOS monochrome camera. Nikon Elements Acquisition Software AR 5.02 was used to collect data. Excitation laser lines used were 405 nm and 488 nm.

#### RNA extraction and quantitative PCR

Total RNA was extracted from cells using the RNeasy Plus Mini Kit (Qiagen, 74134). cDNA was generated from freshly prepared RNA using the iScript cDNA Synthesis Kit (Bio Rad, 1708891) using 250 ng of RNA per 20 µL reaction. Then, master mixes containing 10 µL of Platinum SYBR Green (Thermo Fisher Scientific, 11733038), 7.5 µL of water, and 0.5 µL of 40x primers were made per 20 µL reaction. qPCR reactions were run using a Quantstudio 6 Pro (Thermo Fisher Scientific). The following premixed qPCR primers were obtained from IDT: MIDN (Hs.PT.58.4379975): GCTGCACGAACATCATGT G, CTCAAGGCCGGAACAGTC; IRF4 (Hs.PT.58.2073651): CTTTCCTTTAAACAGTGCCCAAG, GAGAACGAGGAGAAGAGCATC; mTOR (Hs.PT.58.24456023): CCGATTCATGCCCTTCTCTT, CCTTCGTGCCTGTCTGATTC.

#### In Vitro Binding Assays

10 µL of anti-FLAG beads (Sigma, M8823, RRID: AB_2637089) per reaction were rinsed 3 times using 700 µL of CHAPS buffer containing 100 mM NaCl, 40 mM HEPES pH 7.4, 0.5 % CHAPS, and 1x HALT protease/phosphatase inhibitor. After rinsing, the beads were resuspended in 700 µL of CHAPS buffer containing 50 mg/mL of milk (LabScientific, M-0842). The beads were blocked for 1 hour at 4°C with end-to-end rotation. After blocking, the beads were rinsed 3 times with 700 µL of CHAPS buffer. The beads were transferred to low-binding, pre-lubricated Eppendorf tubes (Millipore, CLS3207) and resuspended in 500 µL of CHAPS buffer. 5 µL (19.5 µg per condition) of the indicated FLAG-MsyB-αHelix-C pure protein was added to each tube. The beads were incubated for 1.5 hours at 4°C with end-to-end rotation to immobilize the bait. After incubation, the tubes were rinsed 3 times with 500 µL of CHAPS buffer and resuspended in 500 µL of CHAPS buffer containing the following proteins: TNPO1 (104 kDa) and KPNA2 (63 kDa) were kept equimolar.

Figure 2D: 7.9 µg TNPO1, 4.9 µg KPNA2

Figure 2E: 12 µg proteasome

Figure 2F: 12 µg proteasome and/or 100 µg TNPO1 and/or 62 µg KPNA2

After adding the pure proteins, the beads were incubated end-to-end rotation for 1 hour at 4°C. After incubation, the beads were rinsed three times using 500 µL of CHAPS buffer. The beads were resuspended with 25 µL Tris-Glycine SDS sample buffer and boiled at 95°C for 3 minutes. The purified proteins were collected as input in the same sample buffer. The bait was diluted 1:20 in the same sample buffer for the anti-FLAG blot. 8 µL of each sample was loaded into gels for immunoblotting.

#### Biolayer Interferometry

Stock solutions of proteins were prepared in a buffer containing 50 mM HEPES pH 7.5, 150 mM NaCl, 1 mM DTT, 0.05% CHAPS, and 1% BSA. Specifically, biotinylated FLAG-Avi-MsyB-αHelixC wild-type and 5RD were diluted to a final concentration of 2 µM. The indicated final concentrations of TNPO1 (5, 2, 1, 0.5, 0.2, 0.1, 0.05 µM) and PSMD2 (2, 1, 0.5 µM) were diluted serially from stock solutions. Higher concentrations of PSMD2 were not possible due to yield limitations when purifying PSMD2 from human cells free from the proteasome. We were unsuccessful in purifying PSMD2 from bacteria. Biolayer interferometry measures were made using an Octet RED384 instrument (Sartorius). 60 µL of proteins or buffer were loaded into a 384-well plate. Streptavidin SAX2 biosensors (Sartorius, 18-5136) were first hydrated for 600 seconds in buffer and equilibrated for another 120 seconds in buffer. Biotinylated FLAG-Avi-MsyB-αHelixC wild-type and 5RD proteins were loaded for 120 seconds before a wash step for an additional 120 seconds. Biosensors were then exposed to increasing concentrations of TNPO1 or PSMD2 for 200 seconds of association and then exposed to buffer for 240 seconds of dissociation. BLI data was analyzed using the Octet Analysis Studio software. The dissociation constant (Kd) was determined using the equilibrium approach from the baseline step average, including a baseline step correction and a Savitzky-Golay filter.

#### Mass Spectrometry of 2xFLAG-MBP-αHelix-C

Using lentivirus, *MIDN* knockout HEK-293T cells were reconstituted with 2xFLAG-MBP or 2xFLAG-MBP-αHelix-C variants. Each cell line was grown to >90% confluency in 5 15 cm plates. An anti-FLAG immunoprecipitation was performed as described in the immunoprecipitation section of the materials and methods, using 1 mL of lysis buffer per plate. After the final rinse, the anti-FLAG beads were resuspended with 100 µL of 50 mM Tris pH 8.5 containing 5% SDS, and the protein was eluted off the beads by heating at 95°C for 4 minutes.

The eluted proteins were digested with trypsin on S-Trap Micro columns (Protifi, C02-micro-10). Proteins were first reduced with 5 mM TCEP at 55°C for 15 minutes and alkylated at room temperature with 20 mM iodoacetamide for 30 minutes in the dark. The proteins were then acidified using phosphoric acid to a final concentration of 2.5% (v/v). The protein was then diluted 10-fold with 100 mM Tris, pH 7.55 in 90% methanol/10% water. The solution was passed through S-Trap columns by multiple rounds of centrifugation for 30 seconds at 4000 x g. The trapped protein was then rinsed 3 times with 150 µL of 100 mM Tris, pH 7.55 in 90% methanol/10% water, ending with a dry spin at 4000 x g for 1 minute. A 20 µL solution of 50 mM ammonium bicarbonate, pH 8 containing 2 µg of trypsin was added directly to each column. Digestion was carried out at 37°C in a humid environment overnight. The peptides were eluted from the column by 3 sequential centrifugations for 1 minute at 4,000 x g using first 40 µL ammonium bicarbonate pH 8, second 40 µL 0.2% formic acid in water, and third 40 µL 50% acetonitrile in water. The pooled eluate was dried using a SpeedVac under reduced pressure and the peptides were resuspended in 30 µL 0.1% formic acid in water before data acquisition by LC-MS/MS as previously described ^48^.

A Human UniProt SwissProt proteome was downloaded on December 2^nd^, 2023, and was used as the reference database to search for co-immunoprecipitated proteins. Briefly, FragPipe (v18.0) was used to search the data using the MSFragger search engine. Tryptic peptides with at most two missed cleavages were considered. Fixed peptide modification included carbamidomethylation of cysteine, while oxidation of methionine was a variable modification with at most four variable modifications per peptide. The mass tolerances were 10 ppm for precursor ions and 0.04 Da for product ions. PeptideProphet was used to filter peptide hits to a false discovery rate of 1%.

#### Crosslinking Mass Spectrometry of the Midnolin-Proteasome Complex

Crosslinking reactions were carried out on midnolin-proteasome complexes stored at -80 °C as previously described^59^. Briefly, crosslinking reactions were carried out for 1 hour at room temperature in 100 mM MOPS (piperazine-N,N′-bis(2-ethanesulfonic acid) buffer, pH 6.5, containing 50 mN NaCl, 2 mM MgCl_2_, 75 mM EDC and 16 mM sulfo-*N*-hydroxysuccinimide. Reactions were quenched with hydroxylamine to a final concentration of 75 mM. Samples were reduced for 1 hour in 2% sodium dodecylsulfate and 10 mM tris(2-carboxyethyl)phosphine, alkylated with 20 mM iodoacetamide in the dark for 1 hour and quenched with 20 mM beta-mercaptoethanol for 15 minutes. Samples were then processed using the SP3method^60^ and digested with LysC(Wako) for 3 hours and then trypsin (Promega) for 6 hours, both at 1:25 enzyme:substrate ratio and 37°C. Digested peptides were acidified with 10% TFA to pH ∼2, desalted using stage tips with Empore C18 SPE Extraction Disks (3M), and dried under vacuum.

The sample was reconstituted in 5% FA/5% acetonitrile and analyzed in the Orbitrap Eclipse Mass Spectrometer (Thermo Fisher Scientific) coupled to an EASY-nLC 1200 (Thermo Fisher Scientific) ultra-high pressure liquid chromatography pump, as well as a high-field asymmetric waveform ion mobility spectrometry (FAIMS) interface. Peptides were separated on an in-house pulled 100-μm inner diameter column packed with 35 cm of Accucore C18 resin (2.6 μm, 150 Å, Thermo Fisher Scientific), using a gradient consisting of 5–35% (acetonitrile, 0.125% FA) over 180 minutes at ∼500 nL/min. The instrument was operated in data-dependent mode. FTMS1 spectra were collected at a resolution of 60K, with an AGC target of 100% and a maximum injection time of 50 ms. The most intense ions were selected for tandem MS (MS/MS) for 1.5 s in top-speed mode, while switching among three FAIMS compensation voltages (CVs), −40, –60 and –80 V, in the same method. Precursors were filtered according to charge state (allowed 3 ≤ *z* ≤ 7) and monoisotopic peak assignment was turned on. Previously interrogated precursors were excluded using a dynamic exclusion window (120 s ± 10 ppm). MS2 precursors were isolated with a quadrupole mass filter set to a width of 0.7 *m*/*z* and analyzed by FTMS2, with the Orbitrap operating at 30K resolution, an nAGC target of 250% and a maximum injection time of 150 ms. Precursors were then fragmented by high-energy collision dissociation at 30% normalized collision energy.

Mass spectra were processed and searched using the PIXL search engine^59^. The sequence database contained proteins identified at 1% false discovery rate (FDR) in a noncrosslinked search. For PIXL search^61^, precursor tolerance was set to 15 ppm and fragment ion tolerance to 10 ppm. Methionine oxidation was set as a variable modification. Crosslinked peptides were searched assuming zero-length crosslinker resulting in a loss of water (−18.010565) and considering 80 most abundant protein sequences to ensure sufficient statistics for FDR estimation. Matches were filtered to 1% FDR on the unique peptide level using linear discriminant features as previously described^59^.

#### Midnolin Overexpression and Flow Cytometry

*MIDN* mutant HEK-293T cells, generated previously^5^, were infected with lentivirus to stably overexpress the GPS IRF4 reporter. Once this reporter cell line was established, the cells were seeded in 6-well culture plates at 200,000 cells/well. After growing out the cells for 2 days, the cells were transfected with 1 µg of DNA using PolyJet to overexpress a control blue-fluorescent protein (BFP), or midnolin (wild type or mutant versions) together with BFP under an EF1α promoter. The media was replaced with fresh media one day post-transfection. Two days post-transfection, the cells were detached from the culture plate using 0.05% trypsin and neutralized with fresh media. About 10,000 BFP+ (V-450-PB-A) cells were subsequently analyzed for the GFP/DsRed ratio (B525-FITC-A/Y585-PE-A) on a CytoFLEX S flow cytometer (Beckman Coulter, V2-B2-Y3-R2 version #C09762). The CytExpert software (Beckman Coulter) was used to collect flow cytometry data. Figures were generated using the FlowJo software.

## Acknowledgments

We thank Phil Cole for using his mass spectrometer and J Wade Harper for using his FPLC. SDP acknowledges the technicians Luke Sebastian, Chelsea Braithwaite, Puspalata Bashyal, Coby O’Young, and Kijun Song. Cryo-EM data collection was performed at the Harvard Center for Cryo-Electron Microscopy (HC2EM). Data processing was supported by SBGrid. Biolayer interferometry was performed at the Harvard Center for Macromolecular Interactions. Confocal imaging was performed at the Harvard MicRoN facility. We thank Greenberg, Elledge, and Shao lab members for suggestions and discussions about the project. We thank Tom Rapoport, Dan Finley, Caleb Glassman, and Justine Rutter for their review of the manuscript.

## Author contributions

XG and CN conceptualized this study. CN and XG performed all the biochemical and cellular experiments. MCJY helped establish the protocol for purifying the midnolin-proteasome complex. JG, MCJY, and SS performed cryo-EM sample preparation, data collection, and analysis. HSS and SDP determined the crystal structure of the EGR1-Catch domain. JM and SPG performed the crosslinking mass spectrometry. LO and AKB helped with proliferation assays in myeloma cell lines. MN helped with cloning and designing model diagrams. NK established a platform for AlphaFold multimer predictions. MF and NCM observed midnolin downregulation in multiple myeloma patient data and independently hypothesized a role for midnolin in myeloma growth. XG, SS, SJE, and MEG supervised the study. CN, XG, JG, SS, and SJE wrote the manuscript.

## Supplemental information

Table S2. Mass spectrometry analysis of MBP-αHelix-C co-immunoprecipitation

Table S3. Crosslinking mass spectrometry analysis of the midnolin-proteasome complex

---

**Figure S1.**
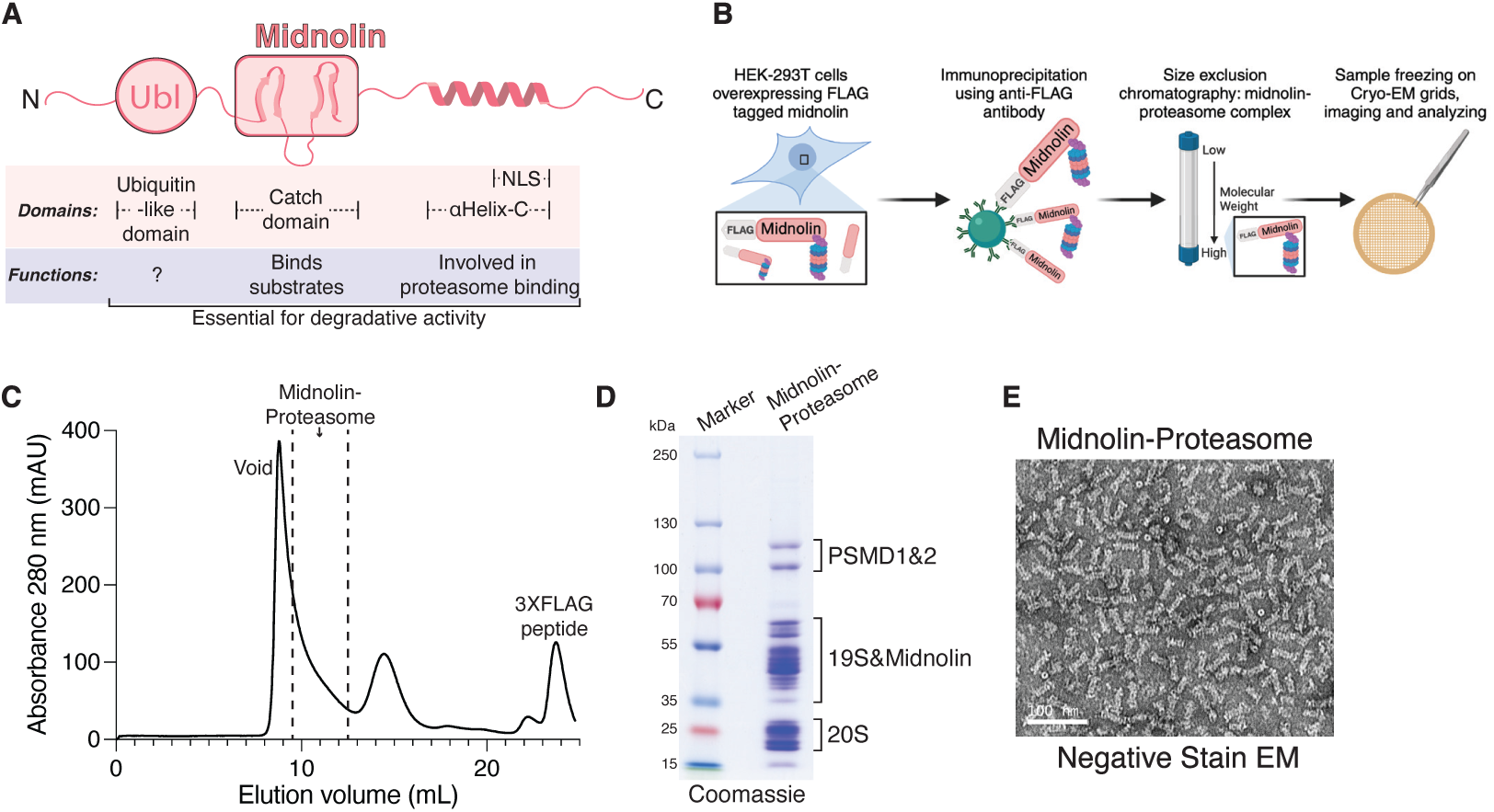
Purification of midnolin-proteasome complexes, related to Figure 1. (A) Midnolin contains three domains necessary for its degradative function: a ubiquitin-like (Ubl) domain, the Catch domain, and a C-terminal helix (αHelix-C) containing a nuclear localization sequence. (B) Schematic representation of the process to affinity-purify the midnolin-proteasome complex by size-exclusion chromatography from HEK-293T cells transiently overexpressing 2xFLAG-tagged midnolin variants. Created with BioRender.com. (C) A representative size-exclusion chromatography trace indicating the fractions collected corresponding to the midnolin-proteasome complex. (D) A representative Coomassie stain of the purified midnolin-proteasome complex showing the characteristic migration pattern of the proteasomal subunits after SDS-PAGE. (E) A representative negative stain electron micrograph of the midnolin-proteasome complex.

**Figure S2.**
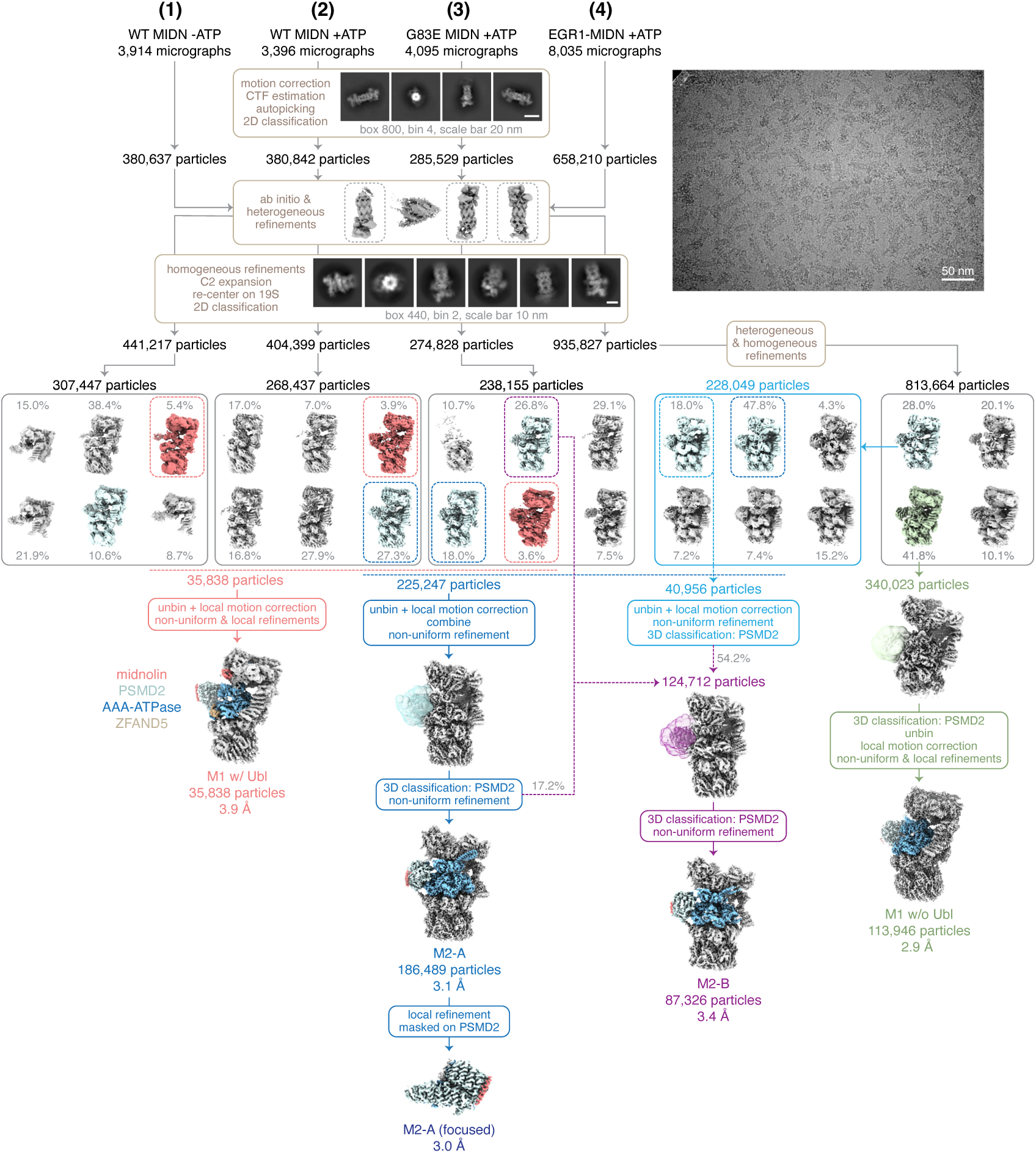
Cryo-EM data processing for the midnolin-proteasome complex, related to Figure 1. Summary of cryo-EM data processing scheme for midnolin-proteasome complexes showing a representative micrograph, 2D classes, and 3D reconstructions. The four datasets derive from four midnolin-proteasome purifications as outlined in the methods.

**Figure S3.**
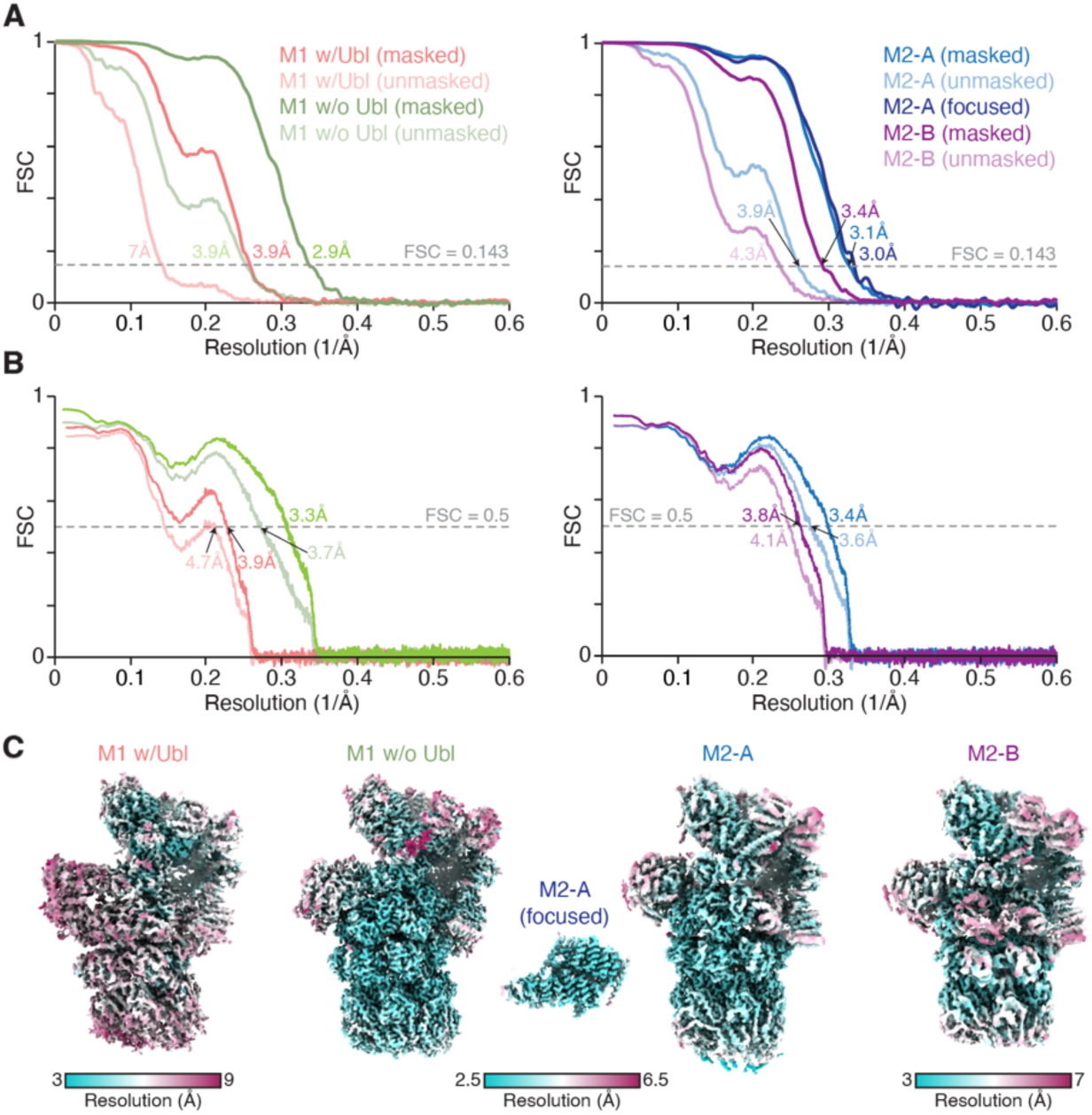
Cryo-EM map and model quality, related to Figure 1. (A) Fourier shell correlation (FSC) vs. resolution (1/Å) curves for the indicated cryo-EM maps. (B) Model vs. map FSC curves. (C) The indicated cryo-EM map is colored by local resolution.

**Figure S4.**
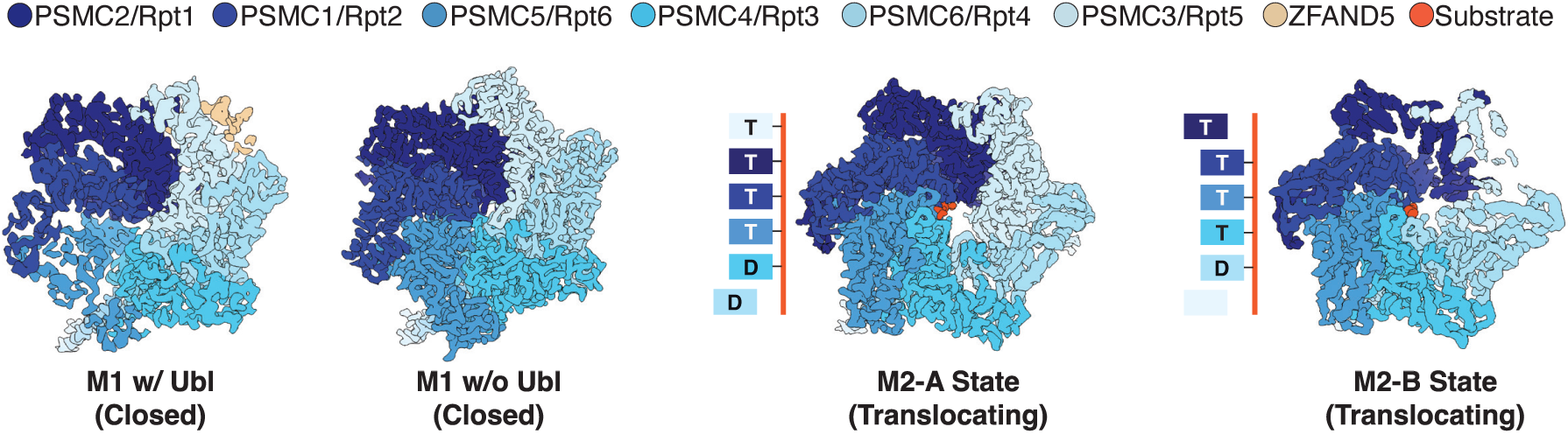
ATPase ring configurations of midnolin-proteasome complexes, related to Figure 1. The density of the ATPase ring observed from the 20S core particle. The central pore of the ATPase ring was closed in the M1 state and open in the M2 state with distinguishable density corresponding to a translocating substrate (orange). A schematic that shows the configuration of the M2 ATPase subunits engaged with a substrate. The nucleotide status of each subunit is indicated by T = ATP, D = ADP, blank = apo.

**Figure S5.**
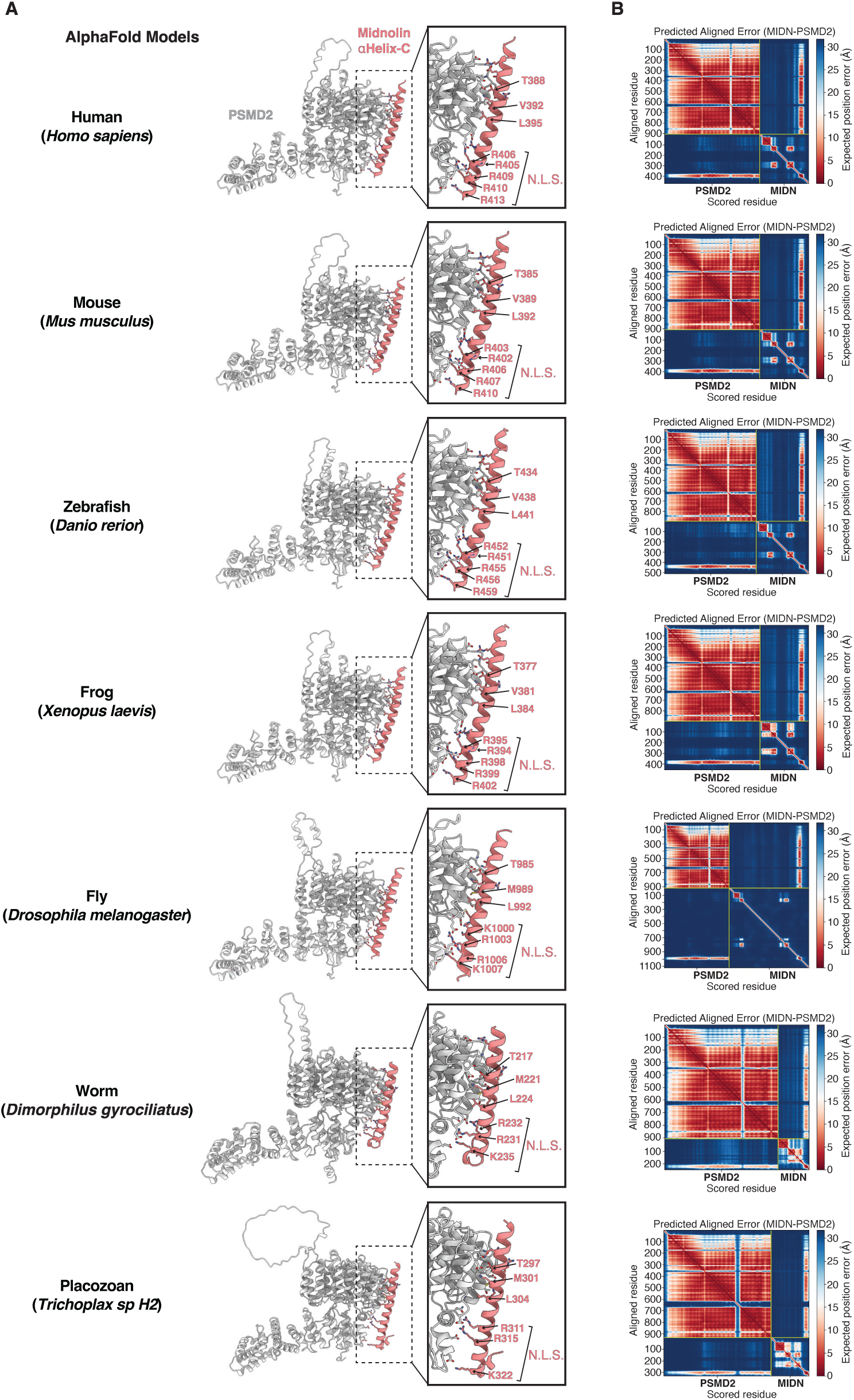
The midnolin-PSMD2 interaction is evolutionarily conserved as predicted by AlphaFold-multimer, related to Figure 2. (A) AlphaFold-multimer predictions of full-length midnolin and PSMD2/Rpn1 using orthologous proteins. (B) The predicted aligned error (PAE) plots of the AlphaFold-multimer predictions suggest the interaction is high confidence.

**Figure S6.**
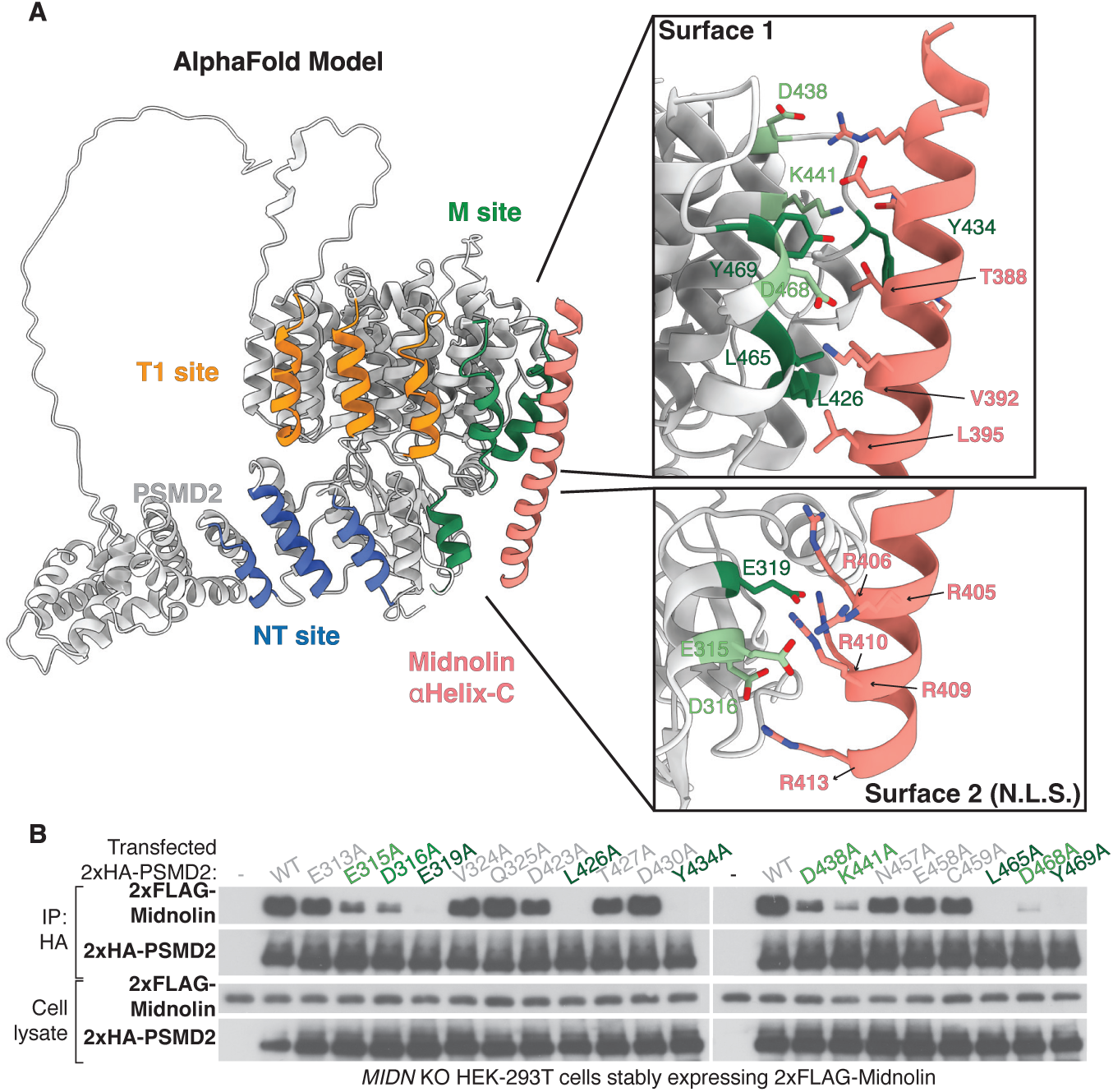
Validating the interaction between αHelix-C and PSMD2, related to Figure 2. (A) AlphaFold-multimer prediction of full-length midnolin with PSMD2/Rpn1. The M site represents PSMD2 residues that make direct contact with αHelix-C. The arginine residues within the midnolin nuclear localization sequence (NLS) mediate a part of the interaction. (B) *MIDN* KO HEK-293T cells were first reconstituted with 2xFLAG-midnolin from a CMV promoter using lentivirus. These cells were then transfected with 2xHA-PSMD2. Shown is immunoblotting from anti-HA immunoprecipitates.

**Figure S7.**
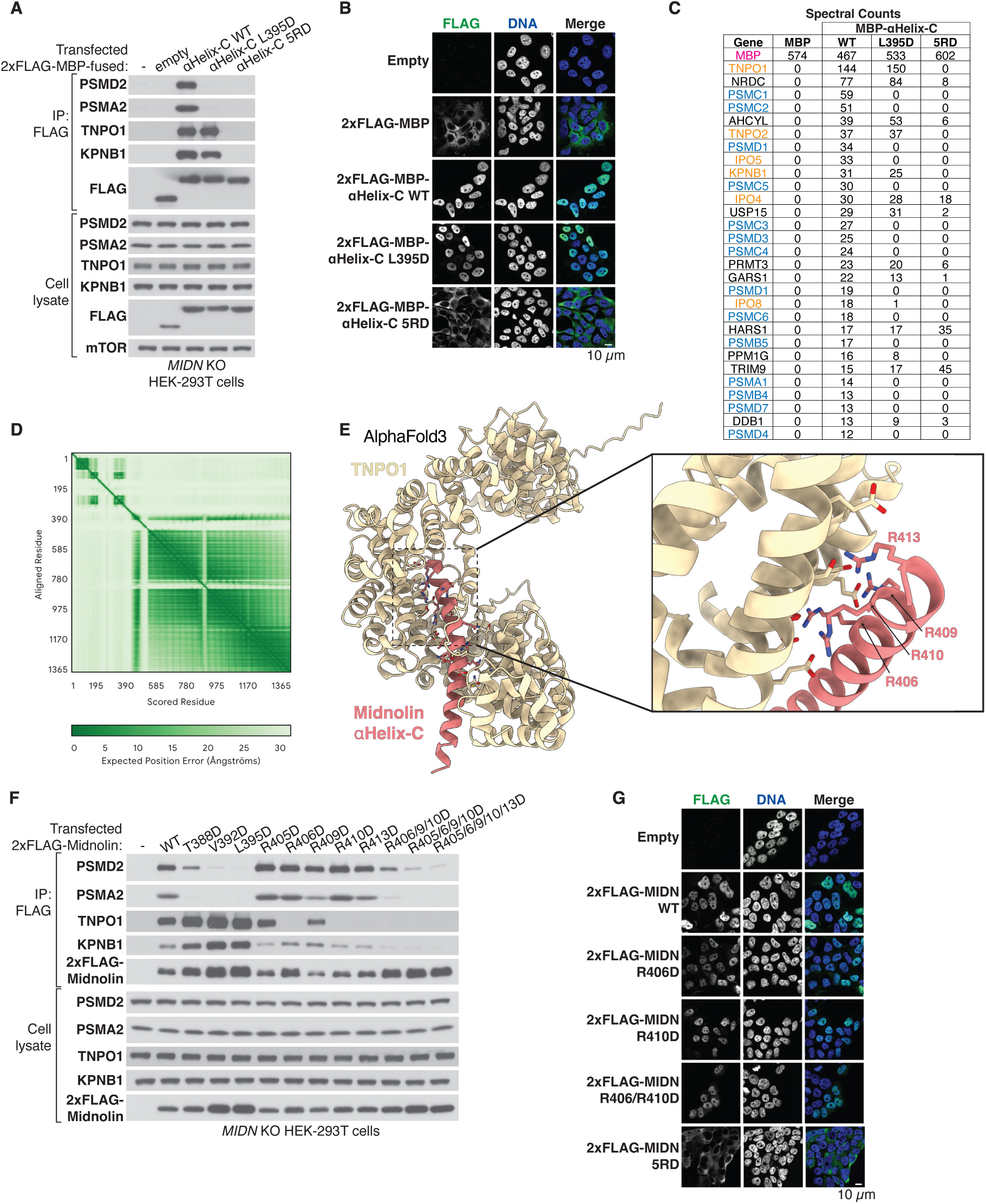
Midnolin binds to the importin β family of nuclear import receptors, related to Figure 2. (A) Immunoblotting of anti-FLAG immunoprecipitants of *MIDN* knockout HEK-293T cells transiently overexpressing 2xFLAG-MBP-αHelix-C variants. (B) Anti-FLAG immunofluorescence of *MIDN* knockout HEK-293T cells stably expressing 2xFLAG-MBP-αHelix-C. (C) Summary of the most enriched co-immunoprecipitated proteins using cell lines from (a) as detected using mass spectrometry. (D) Predicted aligned error (PAE) graph of an (E) AlphaFold3 prediction of full-length midnolin with transportin-1. (F) Immunoblotting was performed from anti-FLAG immunoprecipitants of *MIDN* KO HEK-293T cells that were transiently overexpressing 2xFLAG-tagged midnolin using a CMV promoter. (G) Same assay as (b) but from *MIDN* KO HEK-293T cells stably expressing 2xFLAG-midnolin variants. The cells were treated with 10 µM MG132 for 4 hours.

**Figure S8.**
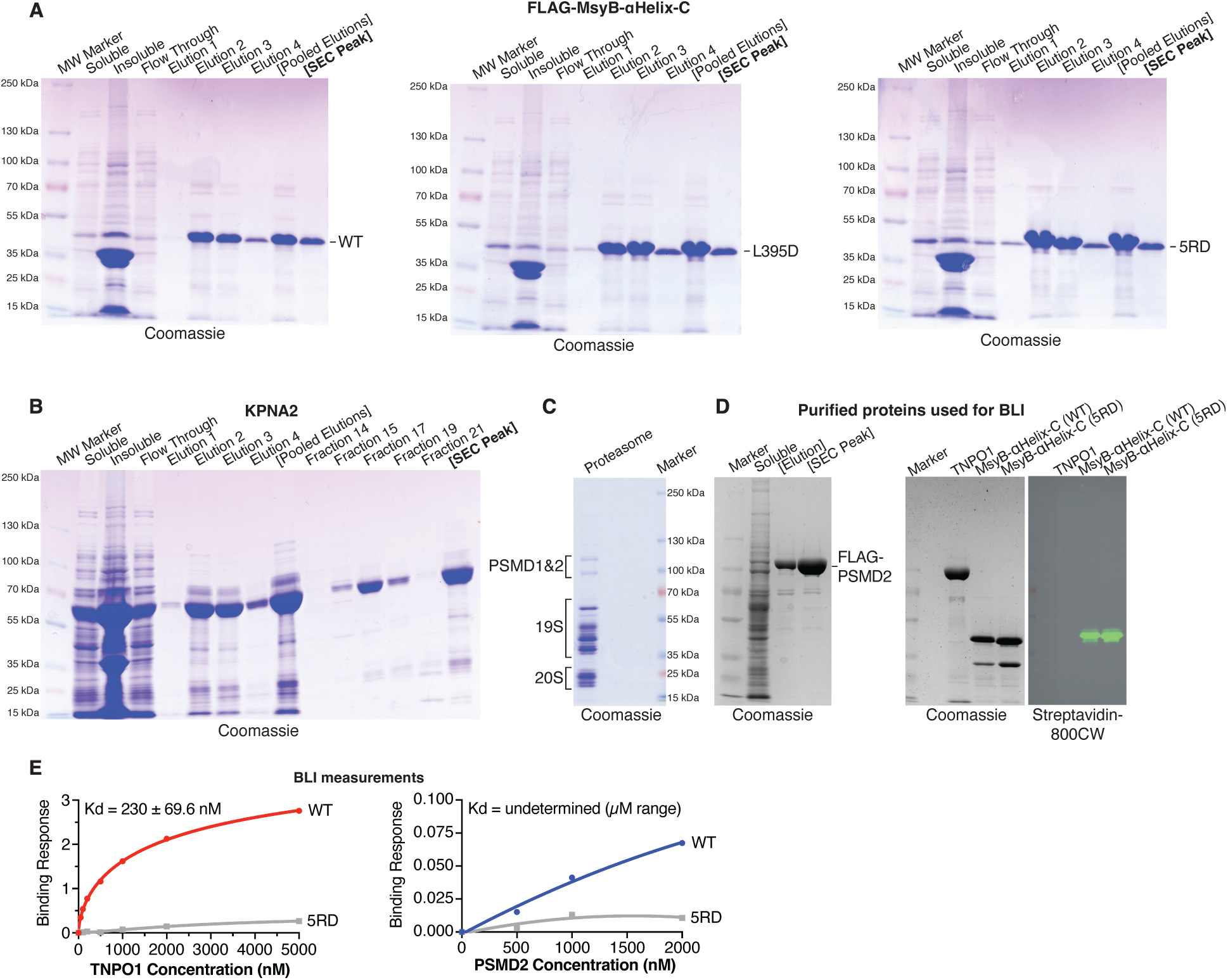
Validations of protein purifications, related to Figure 2. (A) Coomassie stains of recombinant FLAG-MsyB-αHelix-C proteins and (B) recombinant KPNA2 that were purified by size-exclusion chromatography (SEC). The elutions show the 6His-tagged proteins that were first affinity-purified with a Ni-NTA resin before fractionation by size-exclusion chromatography. The concentrated SEC Peak shows the purified protein used in the *in vitro* binding experiments. (C) Coomassie stain of purified human proteasomes showing the characteristic migration pattern of the proteasomal subunits after SDS-PAGE. (D) Coomassie stains of FLAG-PSMD2 purified from human cells, recombinant TNPO1, and recombinant biotinylated Avi-MsyB-αHelix-C variants. The streptavidin-800 is an immunoblot. These proteins were used for biolayer interferometry measurements. (E) Biolayer interferometry measurements using immobilized biotinylated Avi-MsyB-αHelix-C to assess binding with increasing concentrations of TNPO1 or PSMD2.

**Figure S9.**
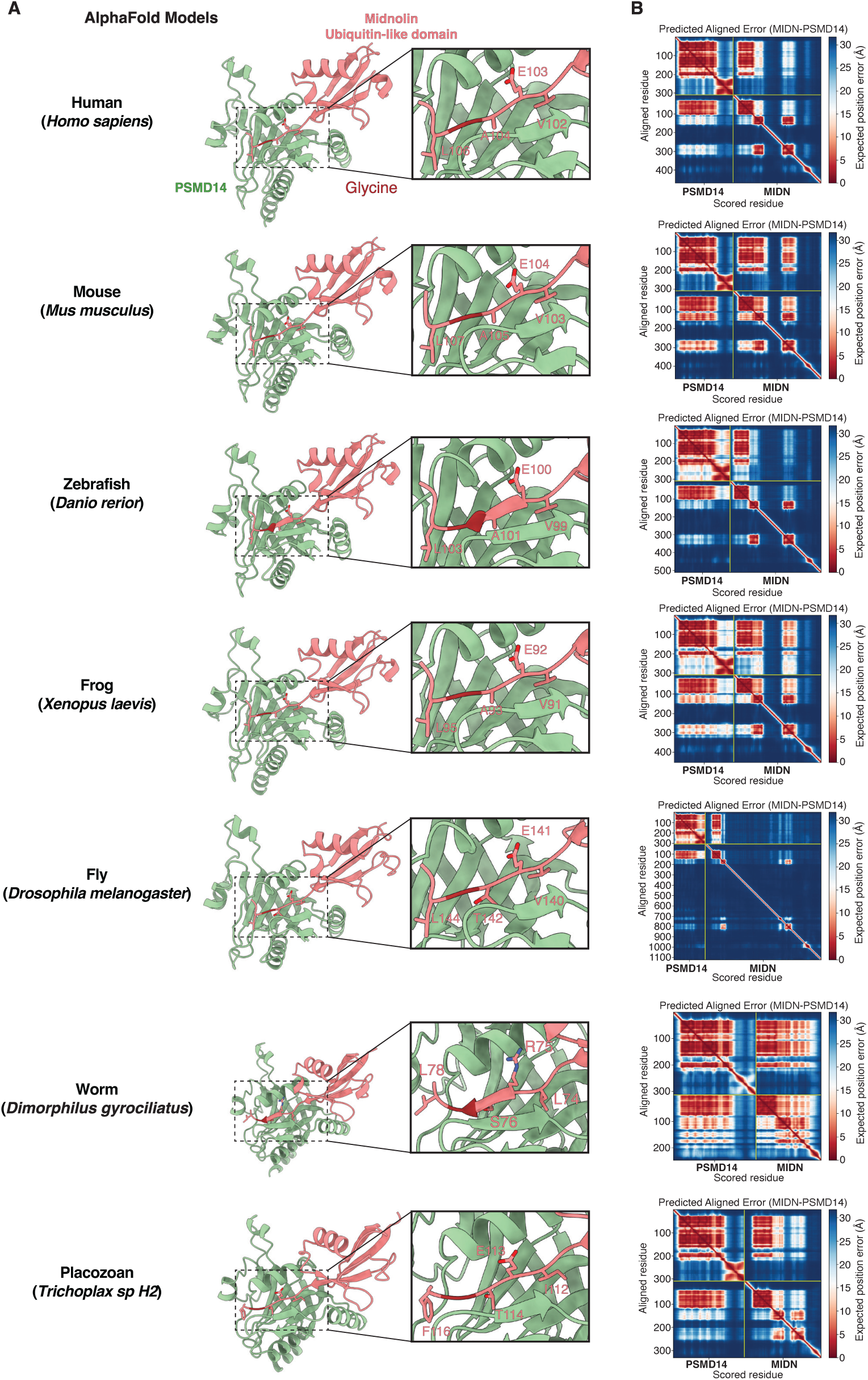
The midnolin-PSMD14 interaction is evolutionarily conserved as predicted by AlphaFold-multimer, related to Figure 3. (A) AlphaFold-multimer predictions of full-length midnolin and PSMD14/Rpn11 using orthologous proteins. (B) The predicted aligned error (PAE) plots of the AlphaFold-multimer predictions suggest the interaction is high confidence.

**Figure S10.**
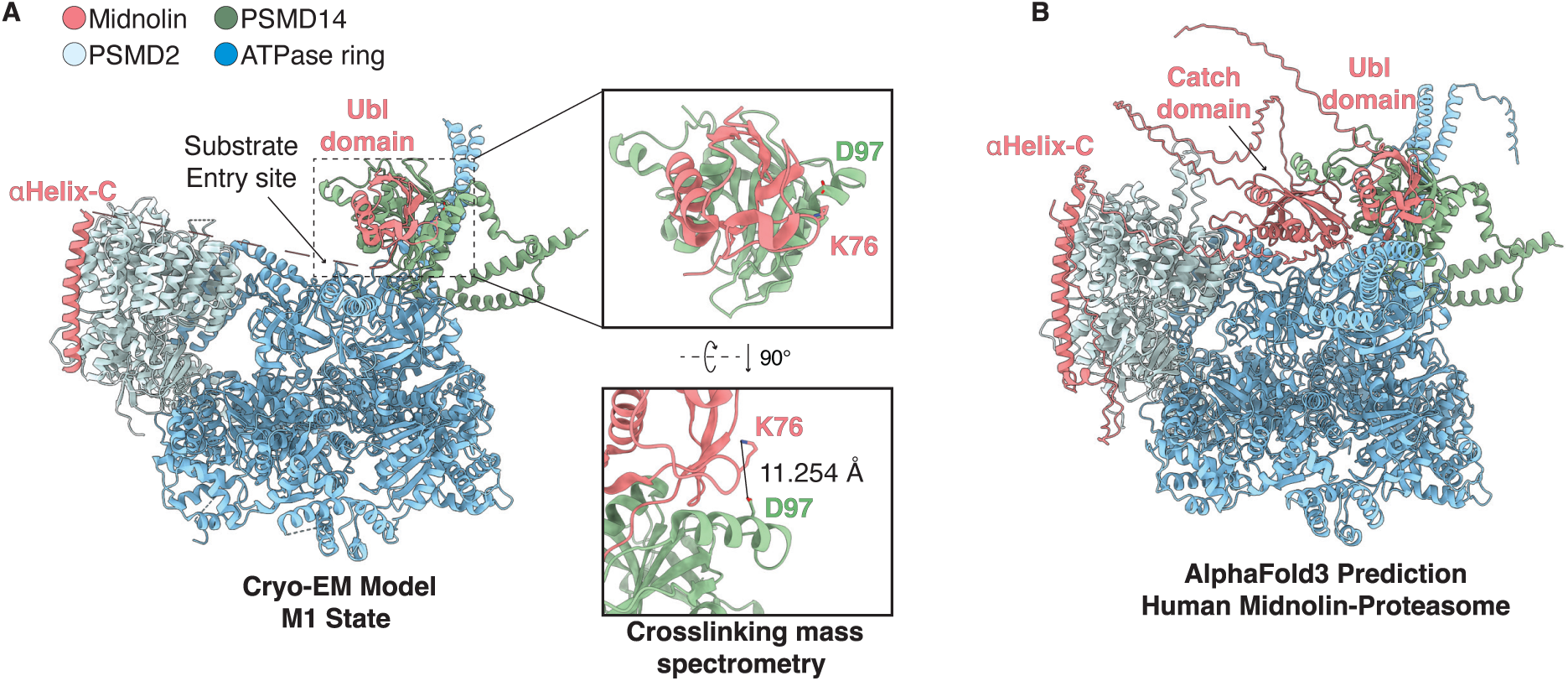
Crosslinking mass spectrometry of the midnolin-proteasome complex, related to Figure 3. (A) Schematic showing the EDC crosslink between the midnolin Ubl domain and PSMD14 detected by mass spectrometry. (B) AlphaFold3 prediction of human midnolin-PSMD2-PSMD14-ATPase subunits resembles the cryo-EM model of the human M1 state. The Catch domain is positioned directly above the ATPase ring.

**Figure S11.**
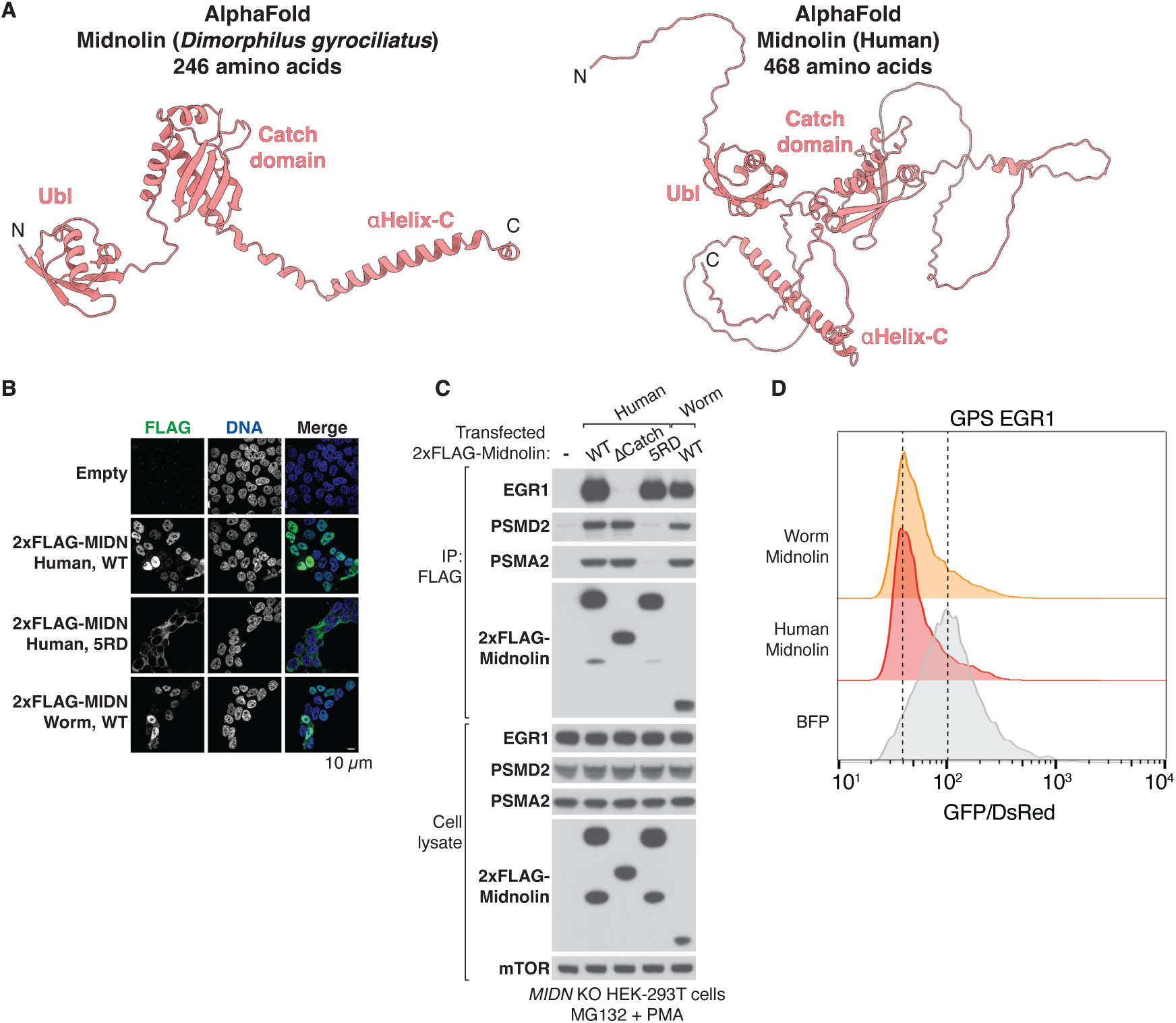
*Dimorphilus gyrociliatus* evolved a miniaturized midnolin that functions in human cells, related to Figure 3. (A) AlphaFold prediction of a minimal midnolin found in *Dimorphilus gyrociliatus* (segmented worm) that contains only the three functional domains: Ubl, Catch, and αHelix-C. (B) anti-FLAG immunofluorescence of *MIDN* KO HEK-293T cells stably expressing 2xFLAG-tagged midnolin variants. Cells were pre-treated with 10 µM MG132 for 4 hours. (C) Immunoblotting from anti-FLAG immunoprecipitations of *MIDN* KO HEK-293T cells transiently overexpressing 2xFLAG-tagged midnolin variants using a CMV promoter. (D) *MIDN* KO HEK-293T cells stably expressing a dual-fluorescence EGR1 stability reporter were transfected with control BFP or midnolin co-expressing BFP using an EF-1α promoter. The BFP+ cells (∼10,000) were analyzed for the GFP/DsRed ratio two days post-transfection by flow cytometry.

**Figure S12.**
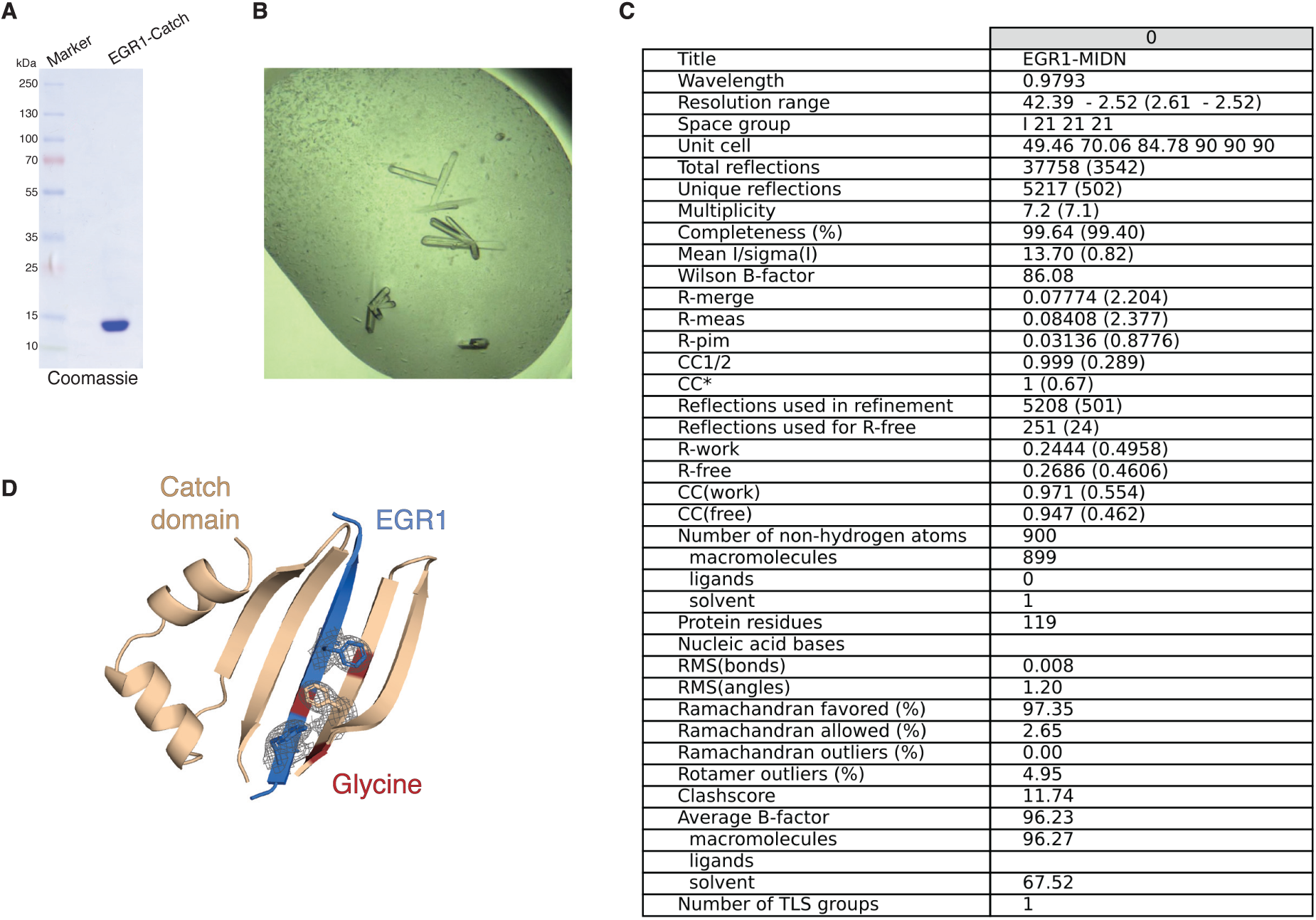
Purification and structure determination of the EGR1-Catch domain fusion, related to Figure 4. (A) Coomassie stain of the recombinant EGR1-Catch fusion protein, which was first affinity enriched and then purified by size-exclusion chromatography. (B) The EGR1-Catch domain crystals harvested for data acquisition. (C) Summary of statistics related to the EGR1-Catch domain crystal structure. (D) The EGR1-Catch crystal structure showing density for the FG zipper residues.

**Figure S13.**
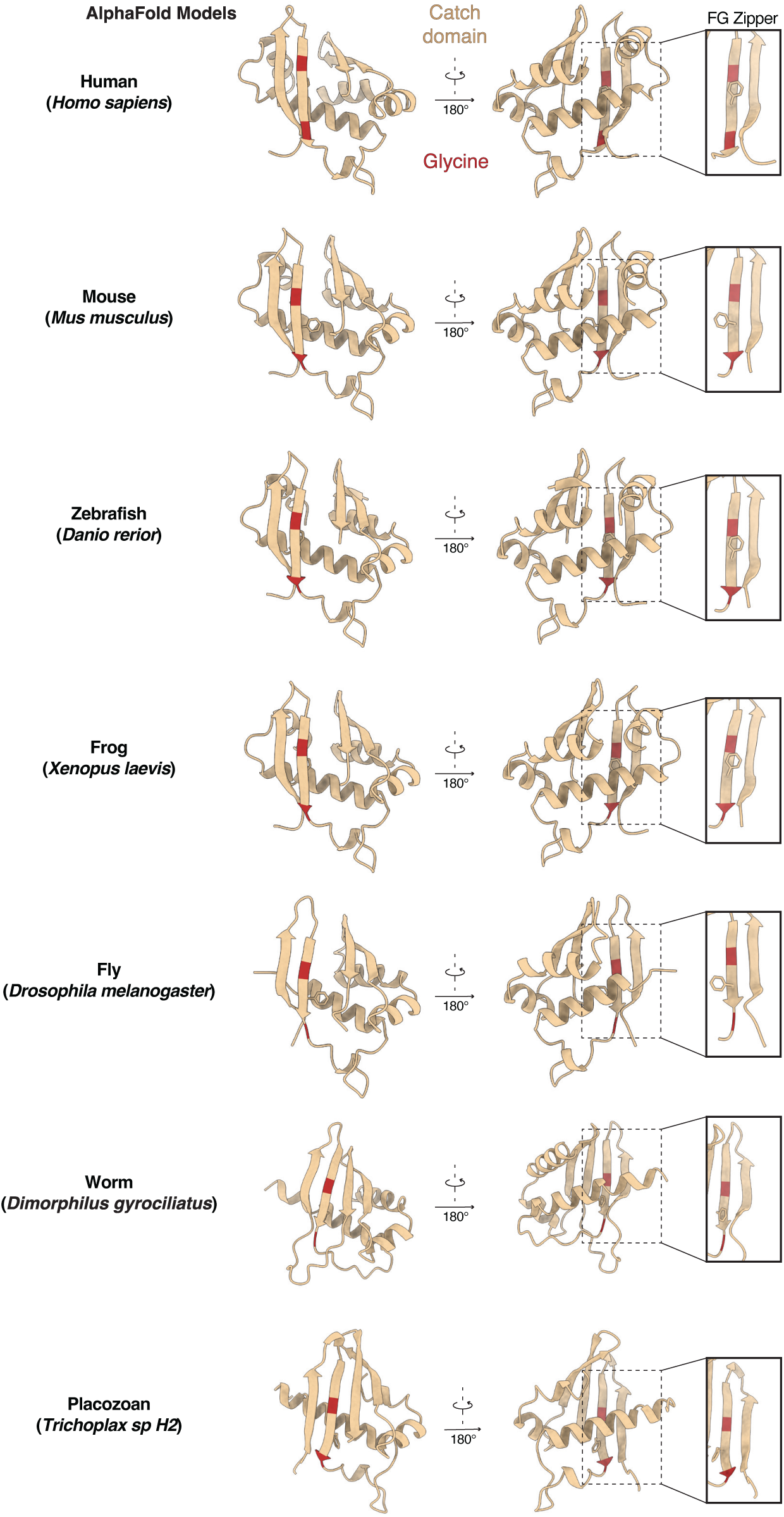
The FG zipper residues of the Catch domain are evolutionarily conserved, related to Figure 4. The Catch domains of orthologous midnolin proteins were extracted from the AlphaFold web server. FG zipper residues and their location within the Catch domain are highly conserved during evolution.

**Figure S14.**
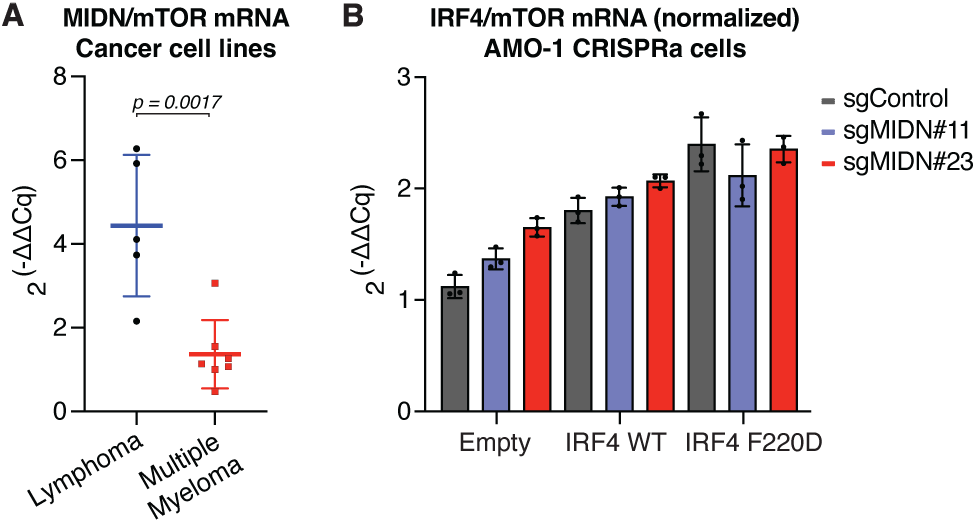
Validation of midnolin downregulation in multiple myeloma, related to Figure 5. (A) qPCR analysis of midnolin and mTOR mRNA from five lymphoma cell lines (BCWM1, RAMOS, MWCL1, SU-DHL-8, Toledo) and seven myeloma cell lines (AMO1, JJN3, KMS27, KMS26, ANBL6, MOLP8, H929). Each data point represents a different cell line with values averaged from technical replicates. Statistical analysis was performed using a two-tailed unpaired *t*-test. (B) qPCR analysis of AMO-1 CRISPR-activation cell lines for IRF4 and mTOR transcripts was performed using biological triplicate.

**Table S1.** Cryo-EM data collection, refinement, and validation statistics, related to Figure 1.

|  | M1 w/Ubl<br>PDB 9BUI<br>EMD-44910 | M2-A<br>PDB 9BV1<br>EMD-44927 | M2-A<br>(focused)<br>EMD-44928 | M2-B<br>PDB 9BV2<br>EMD-44929 | M1 w/o Ubl<br>PDB 9BV3<br>EMD-44930 |
| --- | --- | --- | --- | --- | --- |
| **Data collection and processing** |  |  |  |  |  |
| Magnification | 105,000 | 105,000 |  | 105,000 | 105,000 |
| Voltage (kV) | 300 | 300 |  | 300 | 300 |
| Electron exposure (e-/Å<sup>2</sup>) | 50 to 54 | 50 to 54 |  | 50 to 54 | 54 |
| Defocus range (μm) | -0.8 to -2.2 | -0.8 to -2.2 |  | -0.8 to -2.2 | -0.8 to -1.8 |
| Pixel size (Å) | 0.825 | 0.825 |  | 0.825 | 0.825 |
| Symmetry imposed | C1 | C1 |  | C1 | C1 |
| Initial particle images (no.) | 1,765,798 | 1,765,798 |  | 1,765,798 | 658,210 |
| Final particle images (no.) | 35,838 | 186,489 |  | 87,326 | 113,946 |
| Map resolution (Å) | 3.9 | 3.1 | 3.0 | 3.4 | 2.9 |
| FSC threshold | 0.143 | 0.143 | 0.143 | 0.143 | 0.143 |
| Map resolution range (Å) | 3.4 to 10 | 2.6 to 8.0 | 2.8 to 6.9 | 3.0 to 8.5 | 2.5 to 9.0 |
| **Refinement** |  |  |  |  |  |
| Initial model used | AlphaFold2,<br>ModelAngelo,<br>PDB 7QXW | AlphaFold2,<br>ModelAngelo |  | M2-A,<br>ModelAngelo | M1 w/Ubl,<br>ModelAngelo |
| Model resolution (Å) | 4.4 | 3.4 |  | 3.8 | 3.3 |
| FSC threshold | 0.5 | 0.5 |  | 0.5 | 0.5 |
| Model resolution range (Å) | 4.4 to 62.3 | 3.4 to 59.3 |  | 3.8 to 62.5 | 3.3 to 63.9 |
| Map sharpening *B* factor (Å<sup>2</sup>) | -23.7 | -71.9 | -87.9 | -63.3 | -56.9 |
| **Model composition** |  |  |  |  |  |
| Non-hydrogen atoms | 90,266 | 90,308 |  | 88,606 | 88,575 |
| Protein residues | 11,456 | 11,452 |  | 11,233 | 11,236 |
| Ligands | 2 Mg<sup>2+</sup><br>4 ATP<br>1 ADP | 1 Zn<sup>2+</sup><br>4 Mg<sup>2+</sup><br>4 ATP<br>2 ADP |  | 1 Zn<sup>2+</sup><br>4 ATP<br>1 ADP | 5 Mg<sup>2+</sup><br>4 ATP<br>2 ADP |
| ***B* factors (Å<sup>2</sup>)** |  |  |  |  |  |
| Protein | 208.50 | 70.89 |  | 114.57 | 76.24 |
| Ligand | 215.03 | 50.77 |  | 97.23 | 45.60 |
| **R.m.s. deviations** |  |  |  |  |  |
| Bond lengths (Å) | 0.005 | 0.005 |  | 0.005 | 0.004 |
| Bond angles (°) | 0.989 | 0.981 |  | 0.959 | 0.976 |
| **Validation** |  |  |  |  |  |
| MolProbity score | 2.45 | 2.11 |  | 2.11 | 2.08 |
| Clashscore | 23.07 | 13.23 |  | 13.76 | 12.82 |
| Poor rotamers (%) | 0.01 | 0.02 |  | 0.02 | 0.0 |
| **Ramachandran plot** |  |  |  |  |  |
| Favored (%) | 88.25 | 92.19 |  | 92.68 | 92.79 |
| Allowed (%) | 11.66 | 7.78 |  | 7.30 | 7.18 |
| Disallowed (%) | 0.09 | 0.03 |  | 0.02 | 0.04 |

